# Sex-specific retina–brain signatures link ERα/ERβ imbalance with gliosis in Alzheimer’s disease

**DOI:** 10.64898/2026.04.30.722000

**Authors:** Saba Shahin, Jean-Philippe Vit, Altan Rentsendorj, Dieu-Trang Fuchs, Natalie Swerdlow, Bhakta Prasad Gaire, Edward Robinson, Anna Kissel, Larkin Hagerman, Ayzhah Williams, Alexander V. Ljubimov, Debra Hawes, Lon S. Schneider, Mehdi Mirzaei, Keith L. Black, Yosef Koronyo, Maya Koronyo-Hamaoui

## Abstract

Women face a twofold higher lifetime risk of Alzheimer’s disease (AD) than men, yet the mechanisms underlying female-biased vulnerability and sex-specific disease signatures across the retina-brain axis remain unknown. By integrating clinicopathological and proteomic datasets from paired retinal and brain tissues from 182 donors, we identified sex-divergent molecular and pathological features across the AD continuum. Despite comparable retinal and cerebral amyloid and tau burdens between sexes, females exhibited a more severe neuroinflammatory–neurodegenerative phenotype with intensified gliosis and tissue atrophy, whereas males displayed a dominant vasculopathy, marked by increased retinal vascular Aβ_40_ deposition, tight-junction disruption, and cerebral amyloid angiopathy. In females, this profile coincided with inflammation-associated estrogen receptor (ER)-α remodeling and reduced global and astrocytic-nuclear ER-β, which associated more strongly with cognitive decline than in males. These results indicate that comparable AD proteinopathy is associated with divergent downstream consequences across the retina-brain axis and identify astrocytic ERα/ERβ imbalance as a sex-linked glial mechanism associated with female vulnerability in AD.

## Introduction

Women bear a disproportionate burden of Alzheimer’s disease (AD), accounting for nearly two-thirds of all cases, and facing a two-fold higher lifetime risk (1 in 5) of developing AD compared to age-matched men (1 in 10)^1,2^. Growing evidence shows that women, over the age of 70, experience faster clinical progression and accumulate higher levels of pathological tau in key brain regions essential for memory and cognition, despite comparable cerebral amyloid-beta (Aβ) levels between sexes^3–5^. This disparity is further amplified by the interaction between Aβ, tau, and the apolipoprotein E *ε*4 (*APOEε4*) allele––a major genetic risk factor whose effects are more pronounced in women^6–8^. Moreover, multi-omics studies reveal female-specific dysregulation of key molecular pathways in AD, particularly neuroinflammation, metabolism, and autophagy^9^. Together, these marked sex differences across risk, biomarkers, and disease course emphasize the critical need to define AD mechanisms in women and men for precision diagnostics and treatments.

Mounting evidence indicates that AD-related pathology manifests in the neurosensory retina, a laminated and readily accessible central nervous system (CNS) tissue that shares an embryological origin with the brain^10^. The retina harbors hallmarks AD pathological features, including Aβ-protein deposits, neurofibrillary tangles (NFTs) composed of hyperphosphorylated (p)-tau, gliosis, microvascular dysfunction, and neurodegeneration, in some reports appearing prior to detectable cerebral pathology^10–19^. These features position the retina as a promising, noninvasive imaging site for early detection and longitudinal monitoring of AD progression. Consistent with this, preclinical and human studies suggest that retinal and cerebral AD pathologies follow closely parallel trajectories^13,16,18,20–39^. However, whether this retinal-brain pathological coupling differs by sex and how such differences relate to clinical vulnerability and cognitive decline remains unexplored.

Neuroinflammation is a major contributor to AD pathogenesis and exhibits pronounced sex-specific differences during disease progression^40,41^. Women show heightened cerebral microglial activation and subsequently exacerbated tau pathology and synaptic loss^42^. Furthermore, women exhibit lower basal microglial autophagic activity, impairing the clearance of cerebral Aβ and tau and perpetuating pathological accumulation^43,44^. These sex-specific differences in neuroinflammation are further amplified by the decline in estrogen signaling during menopause, which disrupts neuroprotective pathways mediated by estrogen receptors (ERs): ER-α, ER-β, and G protein–coupled ER1 (GPER1). These receptors are widely expressed in both retina and brain, where they regulate key physiological processes such as inflammation, synaptic plasticity, metabolism, and redox homeostasis^45–51^. Upon ligand binding, ER-α and ER-β translocate to the nucleus, initiating transcriptional programs that enhance synaptic function and neuronal survival, while ER-β also localizes to mitochondria, where it supports oxidative metabolism and neuroprotection. Notably, selective deletion of ER-β in astrocytes, but not neurons, impairs cognitive performance in middle-aged female mice, underscoring its critical role in maintaining cognitive resilience^52^. In AD murine model, estrogen reduces brain Aβ levels by modulating amyloid precursor protein (APP) processing and mitigates tau pathology by enhancing dephosphorylation^53^. Conversely, estrogen loss during menopause disrupts these protective mechanisms, promoting brain Aβ accumulation, tau hyperphosphorylation, and subsequent cognitive decline^54–57^. Although these mechanisms are characterized in the brain, the potential contribution of estrogen signaling dysfunction in the retina, in conjunction with Aβ and tau pathology, neuroinflammation, and vascular alterations, to sex-specific retinal changes has not been previously elucidated.

Here, we investigated whether females and males exhibit distinct retina–brain pathological signatures across early and advanced stages of AD and whether these signatures reflect disease stage and cognitive impairment. We integrated clinical, neuropathological, histopathological, and biochemical/proteome datasets from paired retina and brain tissues across multiple independent cohorts of human donors with normal cognition (NC), mild cognitive impairment (MCI) due to AD, and AD dementia and identified two divergent sex-specific disease trajectories. Despite comparable retina–brain amyloidosis and tauopathy burdens between sexes, AD males exhibited elevated vasculopathy while AD females showed heightened neuroinflammation and neurodegeneration. In females, retinal Aβ_42_ and pathological tau burden were more strongly associated with cognitive impairment and chronic inflammatory responses, whereas in males, pathological burden correlated more closely with disease stage. Additionally, retinal endothelial tight junction protein loss was strongly linked to CAA scores, particularly in males. Proteomic analyses identified estrogen-related signaling as a dysregulated pathway, with ESR1 and ESR2 as emerging upstream regulators in both the retina and brain of AD patients. We further identified astrocytic ER-α upregulation together with an early and marked reduction in global and astrocytic-nuclear ER-β across both retina and brain. Together, our findings define sex-specific AD signatures across the retina–brain axis and identify ERα/ERβ imbalance, particularly in astrocytes, as a molecular axis associated with female-biased gliosis, neurodegeneration, and cognitive vulnerability.

## Results

To investigate sex-specific AD-related retinal histopathological changes and their association with brain pathology and cognitive decline, we analyzed the retina and brain tissues from a well-characterized cohort of 152 individuals. This histological cohort included 92 patients with AD dementia (47 females: mean age ± SD: 82.91±13.58 yrs; 45 males: 78.04±13.30 yrs), 18 individuals with MCI due to AD (MCI; 9 females: 89.56±5.22 yrs, 9 males: 87.78±7.89 yrs), and 42 cognitive normal controls (NC; 24 females: 85.71±12.02 yrs, 18 males: 76.67±9.76 yrs). All analyses were performed using an integrated dataset combining newly generated and previously published immunohistochemical immunoreactivity (IR) or histological measures (total or percent area) of retinal pathological markers quantified from predefined geometrical regions within the superior- and inferior-temporal (ST/IT) retina, alongside brain pathological indices and cognitive scores from our laboratory, derived from female and male individuals spanning normal cognition, MCI due to AD, and AD dementia. Two additional cohorts of 30 individuals were assessed by biochemical methods, including mass spectrometry (MS) and Western blot (WB) analyses, in the temporal retinas (MS: 6AD/6NC; WB: 5-6AD/6NC) and temporal cortices (MS: 10AD/8NC WB: 6AD/6NC). The overall study design is illustrated in Fig. 1a. Importantly, all diagnostic and sex groups were balanced for age, ethnicity, and postmortem interval (PMI) values, ensuring well-matched cohorts for sex-stratified comparisons. Detailed demographics, clinical, and neuropathological characteristics of the cohort are provided in Table 1 and Table S1-3. In the histological cohort, both female and male MCI and AD patients compared to their respective NC controls exhibited significantly elevated Clinical Dementia Rating (CDR) scores (Fig. 1b, female: MCI vs NC = 11.6-fold, *P < 0.0001;* AD vs NC = 14.8-fold, *P < 0.0001;* male: MCI and AD = 5.7-fold, *P = 0.006*; AD vs NC=6.9-fold, *P < 0.0001*) and reduced Mini-Mental State Examination (MMSE) scores (Fig. 1c, female: AD vs NC = 56.5%, *P < 0.0001,* MCI vs AD = 40%, *P = 0.01;* male: AD vs NC = 60%, *P < 0.0001,* MCI vs AD = 47%, *P = 0.001*). Whereas mild sex-related differences were apparent in CDR and MMSE trajectories, with AD females showing slightly higher CDR scores than AD males and MCI males exhibiting a modestly higher MMSE than MCI females, these trends did not reach statistical significance (Fig. 1b,c).

**Fig. 1.**
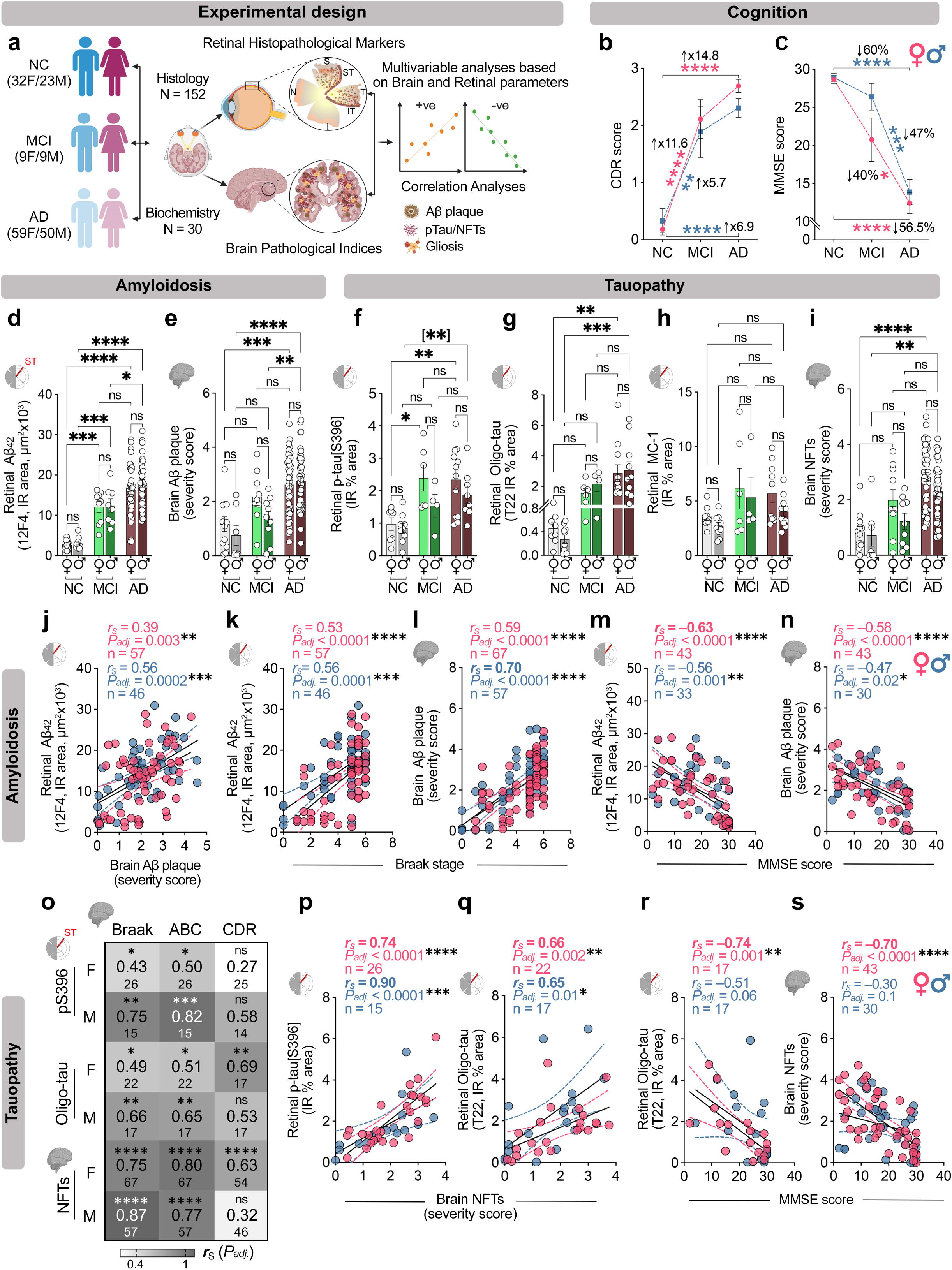
Sex-dependent interactions between amyloidosis and tauopathy across the retina and brain with disease status. **a** Experimental design. Retinal and cerebral tissues from a well-characterized cohort of 182 individuals were analyzed. Histological cohort included 91 patients with AD dementia (47 females; 45 males), 18 individuals with mild cognitive impairment due to AD (MCI; 9 females, 9 males), and 42 cognitively normal controls (NC; 24 females, 18 males). Analyses were performed after combining newly generated and previously published immunohistochemical immunoreactivity (IR) or histological measures (total or percent area) of retinal pathological markers quantified from predefined geometrical regions within the superior-and inferior-temporal (ST/IT) retina (red strips) and correlated with brain pathological indices from female and male MCI, AD, and NC individuals. Two additional cohorts were assessed by biochemical methods, including mass spectrometry (MS) and Western blot (WB) analyses, in the temporal retinas (n=6AD/6NC donors) and temporal cortices (n=18MS+10WB=28). **b-c** Clinical dementia rating (CDR, **b;** NC=11F/6M, MCI= 9F/9M, AD= 34F/31M) and mini-mental state examination (MMSE, **c;** NC=12F/12M, MCI=8F/5M, AD=26F/19M) scores in a subset of individuals stratified by sex and diagnosis. **d,e** Retinal and cerebral Aβ burdens in a subset of age-matched female and male individuals with NC (n=12-17F/7-12M), MCI (n=9F/8-9M) and AD dementia (n= 38-46F/36-41M). **f-h** Retinal tau pathology quantified by measuring the IR % area for **(f)** p-Tau[S396] (NC=7F/9M, MCI= 6F/4M, AD= 13F/8M), **(g)** T22^+^ Oligomeric (Oligo)-tau (NC=8F/10M, MCI= 6F/4M, AD= 11F/10M), and **(h)** MC-1^+^ pre- and mature-tangles; NC=8F/9M, MCI= 6F/4M, AD= 10F/10M) in a subset of age-matched female and male individuals stratified by clinical diagnosis. **i** Brain neurofibrillary tangles (NFTs) severity score in female and male individuals with NC (n=12F/7M), MCI (n=9F/9M) and AD (n= 45F/39M). Data shown as individual subjects (circles) with group means ± SEMs. Statistical analyses used 2-way ANOVA with Bonferroni’s post hoc tests for 3- or more group comparisons and 2-tailed unpaired Student’s *t*-test for 2-group comparisons (in parentheses). **j** Spearman correlation between retinal Aβ_42_ IR area and brain Aβ plaque severity score, **k-n** Spearman correlation between retinal Aβ_42_ IR area or brain Aβ plaque and **(k,l)** Braak stage, and **(m,n)** MMSE cognitive scores. **o** Heatmap showing Spearman’s correlations (*r_S_*) and significance levels (asterisks) between retinal tau species or brain NFTs and brain AD pathology (BRAAK, ABC scores) and CDR score. **p-s** Spearman correlation coefficient analyses between retinal p-tau[S396] or Oligo-tau IR % area and **(p,q)** brain NFTs severity score and **(r,s)** MMSE scores. Correlation coefficient value, Holm-Šídák adjusted p-values (asterisks) and patient number (n) are shown in the upper right corner of each graph. *\*P ≤ 0.05, **P ≤ 0.01, ***P ≤ 0.001, ****P ≤ 0.0001, ns: nonsignificant*.

**Table 1.** Demographic and neuropathological data of female and male human donors used for histology.

| Diagnosis |  | NC | MCI | AD | *F* | *P* |
| --- | --- | --- | --- | --- | --- | --- |
| **Females + Males** |  |  |  |  |  |  |
| *n* = 152 |  | 42 | 18 | 92 | — | — |
| Age at death (years) |  | 81.8 ± 11.9 | 88.7 ± 6.5 | 80.6 ± 13.6 | 3.18 | **0.044** |
| PMI (hours) |  | 10.1 ± 7.9 | 11.8 ± 10.8 | 12.0 ± 12.2 | 0.41 | 0.66 |
| **Females** |  |  |  |  |  |  |
| *n* = 80 |  | 24 | 9 | 47 | — | — |
| Age at death (years) |  | 85.7 ± 12.0 | 89.6 ± 5.2 | 82.9 ± 13.6 | 1.2 | 0.30 |
| Race |  | 22W (92%),<br>1B, 1H | 8W (89%),<br>1B | 28W (62%),<br>4B, 6A, 7H | — | — |
| PMI (hours) |  | 11.4 ± 9.5 | 8.4 ± 6.1 | 11.7 ± 11.0 | 0.41 | 0.66 |
| Brain neuropathology<br>(*n* = 67) | Braak stage | 2.58 ± 1.22 | 3.89 ± 1.65 | 6.34 ± 7.38 | 32.72 | **<0.0001** |
|  | Stage 0-II | 50% | 22% | 2% |  |  |
|  | Stage III-IV | 42% | 33% | 9% |  |  |
|  | Stage V-VI | 8% | 44% | 89% |  |  |
|  | ABC score | 1.65 ± 0.79 | 2.33 ± 0.50 | 2.83 ± 0.31 | 32.74 | **<0.0001** |
|  | NT severity score | 1.12 ± 0.95 | 1.49 ± 1.43 | 2.14 ± 1.51 | 2.84 | 0.066 |
| **Males** |  |  |  |  |  |  |
| *n* = 72 |  | 18 | 9 | 45 | — | — |
| Age at death (years) |  | 76.7 ± 9.8 | 87.8 ± 7.9 | 78.0 ± 13.3 | 2.9 | 0.062 |
| Race |  | 13W (72%),<br>2B, 3H | 8W (89%),<br>1H | 32W (73%),<br>2B, 3A, 5H | — | — |
| PMI (hours) |  | 8.0 ± 3.6 | 15.2 ± 13.6 | 12.3 ± 13.5 | 1.08 | 0.35 |
| Brain neuropathology<br>(*n* = 57) | Braak stage | 1.21 ± 1.78 | 2.78 ± 1.62 | 4.88 ± 1.05 | 31.82 | **<0.0001** |
|  | Stage 0-II | 86% | 44% | 2% |  |  |
|  | Stage III-IV | 0% | 33% | 22% |  |  |
|  | Stage V-VI | 14% | 22% | 76% |  |  |
|  | ABC score | 1.24 ± 0.79 | 1.89 ± 0.69 | 2.80 ± 0.33 | 38.93 | **<0.0001** |
|  | NT severity score | 0.64 ± 1.38 | 0.87 ± 0.76 | 2.27 ± 1.56 | 6.09 | **0.0042** |

Paired brains with neuropathological assessments were available for 124 human donors. Diagnosis: NC, cognitively normal controls; MCI, mild cognitive impairment; AD, Alzheimer's disease. Race: A, Asian; B, Black; H, Hispanic; W, White. PMI, postmortem interval. ABC score: A, Aβ plaque score modified from Thal; B, NFT stage modified from Braak; C, neuritic plaque score modified from CERAD. NT, neuropil threads. Group values are presented as mean ± standard deviation. *F* and *P*-values were determined using one-way ANOVA.

### Retinal and cerebral Aβ/tau pathology does not differ by sex but links to cognition preferentially in females

To determine sex-specific retinal and brain amyloidosis levels, we quantified and compared the retinal 12F4^+^ Aβ_42_ IR area and brain Aβ-plaque severity score in female versus male donors across MCI, AD, and NC controls. In females, we find that retinal Aβ_42_ IR area is significantly increased in both MCI (4.5-fold, *P = 0.0001*) and AD (6.1-fold, *P < 0.0001*) compared to NC controls (Fig. 1d). Similarly, relative to NC males, significantly elevated retinal Aβ_42_ IR area is observed in MCI males (4.2-fold, *P = 0.0008*) and AD (6-fold, *P < 0.0001*). Within the AD group, both sexes showed 1.4-fold higher retinal Aβ_42_ burden compared to their respective MCI counterparts, however, this increase reached statistical significance only in males (Fig. 1d, *P* = 0.0008). Importantly, no sex differences in retinal Aβ_42_ levels were detected between females and males within each diagnostic group. These findings mirror matched brain tissues, where Aβ plaque burden increased significantly with disease progression but did not differ significantly by sex within diagnostic groups (Fig. 1e). Similar sex-independent accumulation was observed when retinal and brain amyloidosis stratified per Braak stage, ABC, CDR, and MMSE scores, and *APOEε4* genotype (Extended Data Fig. 1a-f). Interestingly, Braak stage-based stratification revealed a mid-stage specific divergence at Braak III–IV, where males exhibited significantly higher Aβ burden than females, with a 1.6-fold increase in the retina and a 1.7-fold increase in the brain (Extended Data Fig. 1a, d; *Stage III-IV; P = 0.01-0.02*, n = 12-13F/12-38M donors). Together, these data indicate that retinal and cerebral amyloidosis largely track clinical and neuropathological progression in a sex-independent manner.

Next, we examined sex-specific differences in retinal tau burden by quantifying multiple pathological tau species, including p-tau[S396], oligomeric tau (Oligo-tau; T22), and pretangle/mature tau tangles (MC-1), in postmortem retinas from female and male MCI and AD patients. Retinal p-tau[S396] IR area (%) was significantly increased in both females and males, showing 1.9-fold elevations in MCI and a 2.1–2.2-fold increases in AD relative to sex-respective NC controls (Fig. 1f). Similarly, retinal oligomeric tau burden was significantly higher in both female and male AD patient retinas compared to their respective NC controls (Fig. 1g). A non-significant higher trend in pretangle/mature tangle pathology (MC-1 immunoreactivity) was also observed in the retinas of both MCI and AD individuals, independent of sex (Fig. 1h). The tendency toward greater retinal tauopathy in females than in males mirrored the corresponding trend in cerebral NFT burden (Fig. 1f–i). Retinal p-tau and brain NFT levels, stratified by Braak stage and ABC score categories, CDR or MMSE cognitive status, and *APOEε4* allele carriers, further revealed consistent trends of increases in tauopathy burden in females versus males at different disease stages (Extended Data Fig. 2a-f).

We next examined sex-dependent associations between retinal and cerebral amyloid and tau pathology and AD-related neuropathologic burden, disease stage, and cognitive performance. Spearman correlations revealed stronger correlations in males versus females between retinal Aβ_42_ burden and brain Aβ-plaque severity (Fig. 1j; *r_S_* = 0.39–0.56; *P_adj_ ≤ 0.003–0.0002*). In contrast, comparable strength of correlations was observed in females versus males for retinal Aβ_42_ or brain Aβ-plaque burdens against Braak stage or ABC scores (Fig. 1k-l and Extended Data Fig. 3a,b; female: *r_S_* = 0.51–0.70; *P_adj_ < 0.0001*; male: *r_S_* = 0.56-0.70; *P_adj_ < 0.001–0.0001*).

Notably, the relationship between retinal Aβ_42_ and MMSE or CDR cognitive performance was markedly stronger in females (Fig. 1m and Extended Data Fig. 3c; *r_S_* = 0.61 to –0.63, *P_adj_ < 0.0001*), whereas in males these associations were only moderate (*r_S_* = 0.41 to –0.56, *P_adj_ = 0.008–0.001*). These retinal amyloidosis findings paralleled our observations in the brain, where cerebral Aβ correlates moderately but more strongly in females versus males with MMSE and CDR cognitive performance (Fig. 1n and Extended Data Fig. 3d; female: *r_S_* = 0.50 to –0.58, *P_adj_ < 0.0001*; male: *r_S_* = 0.34 to –0.47, *P_adj_ = 0.02*). These data suggest that retinal Aβ tracks cognitive status more closely than brain Aβ, with associations consistently stronger in females than in males.

Assessing the relationship between retinal tau species and disease status further uncovered sex-specific differences. In males, retinal p-tau[S396] and oligomeric tau burdens showed strong to very-strong correlations with Braak stage, ABC score, and brain NFTs, whereas in females these were moderate to strong (Fig. 1o,p; male: *r_S_* = 0.65–0.90; *P_adj_ < 0.05–0.0001*; female: *r_S_* = 0.43–0.74; *P_adj_ < 0.05–0.0001*). Brain NFT burden correlated strongly to very strongly with Braak stage and ABC scores in both sexes (Fig. 1o, *r* = 0.75-0.87, *P_adj_ < 0.0001*). Importantly, in females, retinal oligomeric tau and brain NFT burdens strongly linked to cognitive status (Fig. 1o,r,s; CDR: *r_S_* = 0.63–0.69 and MMSE: *r_S_* = –0.70 to –0.74; *P_adj_ < 0.01–0.0001*), whereas in males, associations with cognition did not reaching statistically significant. Retinal MC1^+^ tau tangles showed no significant associations with brain pathological indices or cognitive status, and retinal p-tau[S396] was not significantly associated with cognitive measures, in either sex (Extended Data Fig. 3e-j). Consistent with these patterns, retinal oligomeric tau correlated strongly with retinal Aβ_42,_ similar to brain NFTs with cerebral Aβ plaque burden in both sexes, with a significantly stronger association in females (Extended Data Fig. 3k-n), suggesting that amyloid accumulation may be more tightly associated with tau oligomerization in females than in males.

Together, these findings indicate that retinal and brain Aβ/tau pathologies track disease severity and stage in both sexes, yet relate more tightly to cognitive impairment in females, revealing a sex-specific coupling between tissue pathology and clinical outcomes.

### Sex-specific neurodegenerative and vascular vulnerability across the retina–brain axis

Given that females with AD typically exhibit a more severe and accelerated neurodegenerative trajectory than males, particularly at advanced stages^4,58^, we asked whether this sex bias extends to the retina by quantifying neurodegenerative signatures in age-matched female and male retinas and brains across NC, MCI and AD dementia. Retinal neurodegeneration was assessed by total retinal thickness and Nissl^+^ neuronal area, as an integrated measure of neuronal loss, and cerebral neurodegeneration by brain atrophy severity score (Fig. 2a,b and Extended Data Fig. 4a). Retinal thinning and brain atrophy increased with disease stage in both sexes. At the MCI stage, females and males showed comparable neurodegeneration in the retina and brain, displaying significant reductions in retinal thickness/neuronal area and a trend toward exacerbated brain atrophy versus NC, with no significant sex differences. By AD dementia, however, females exhibited greater neurodegeneration across the retina–brain axis, with thinner retinas than males (10% reduction, *P = 0.0308*; n = 33F/29M) and greater brain atrophy (1.2-fold, *P = 0.045*; n = 46F/41M). Consistent with these sex-dependent neurodegenerative trajectories, retinal thickness was moderately and inversely associated with retinal Aβ_42_ burden in females (Fig. 2c, *r_P_* = –0.55; *P_adj_ = 0.00004*; n = 55), whereas in males, there was no significant association (*r_P_* = –0.22; *P_adj_ = 0.4*; n = 49), implicating enhanced retinal vulnerability to Aβ_42_ accumulation in females. No associations were found between retinal thickness and retinal p-tau[S396] or T22^+^ oligomeric tau (Fig. 2d and Extended Data Fig. 4b; *r_P_* = –0.13-0.26; *P_adj_ = 0.4-0.8*; n = 22-24F/19-21M). Brain atrophy showed only weak trends with cerebral Aβ and NFT burden in females and no detectable associations in males (Fig. 2e,f; *r_S_* = 0.006–0.23; *P_adj_ = 1.0–0.06*; n = 57–67F/57). Retinal thickness and brain atrophy did not differ by *APOEε4* status (Fig. 2g). In females, retinal thickness correlated weakly but significantly with brain atrophy severity (Fig. 2h; *r_S_* = –0.28, *P_adj_ = 0.006*, n = 53), and both retinal thickness and brain atrophy showed weak-to-moderate associations with Braak stage and ABC score (Fig. 2i–j and Extended Data Fig. 4c,d; *r_S_* = –0.27 to 0.42; *P_adj_ = 0.049–0.0008*; n = 52–67); these relationships were not significant in males. Consistently, stratification by Braak stage and ABC score indicated more progressive neurodegeneration in females at advanced stages (Extended Data Fig. 4e–h). Associations with cognition also diverged across tissues: retinal thickness associated with CDR in females but not males, whereas brain atrophy showed associations with cognitive status (CDR, MMSE) in both males and females (Fig. 2k,l and Extended Data Fig. 4i,j; *r_S_* = –0.20–0.50; *P_adj_ = 0.19–0.0007*; n = 41–54F/26–46M). Together, these data indicate comparable early retinal neurodegeneration across sexes, but greater female vulnerability to advanced brain–retina atrophy, with female-biased associations with pathological stage and cognition that were weak to moderate in magnitude.

**Fig. 2.**
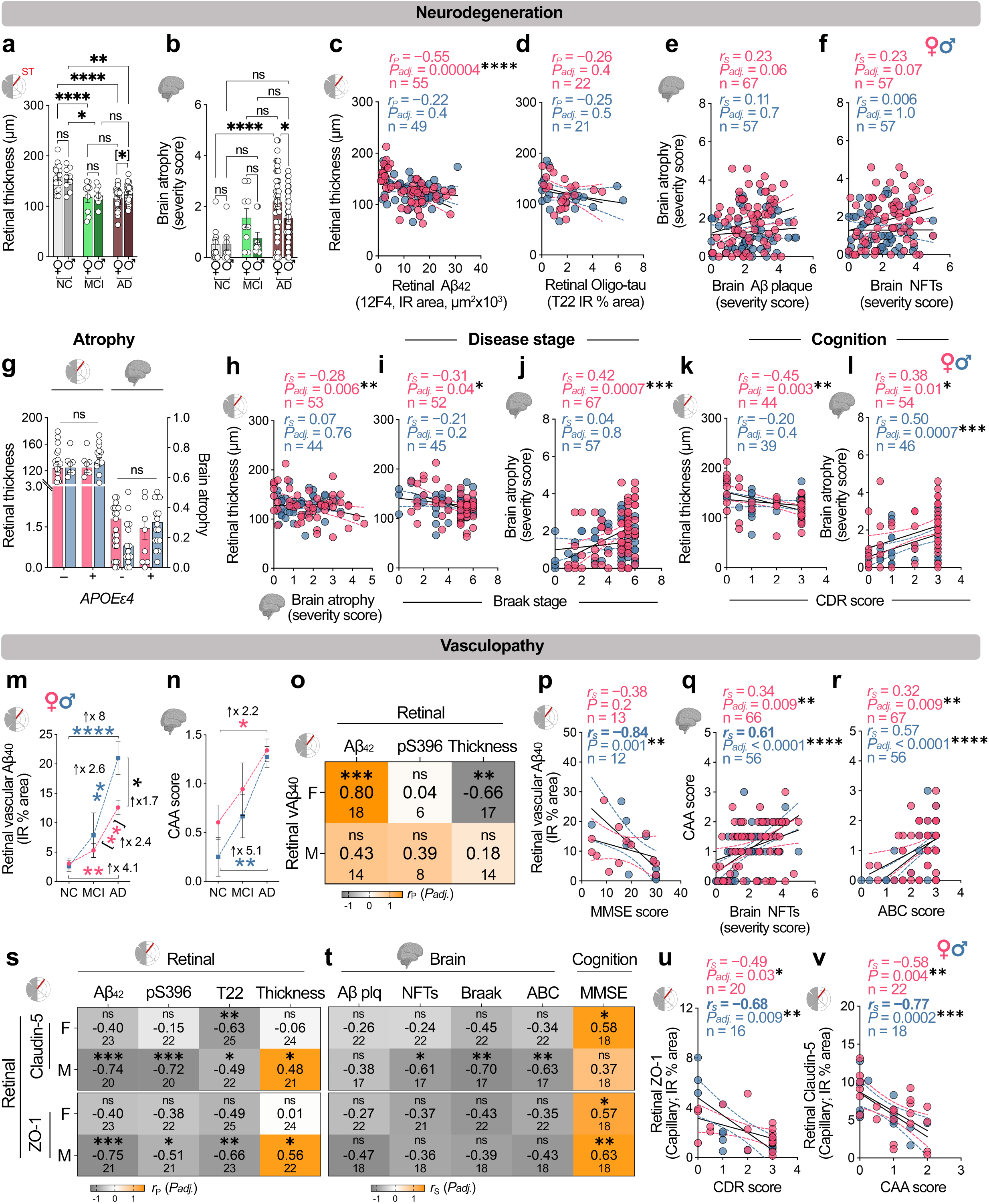
Retina–brain neurodegeneration and vasculopathy are coupled to disease severity and differ by sex. **a** Retinal thickness, a surrogate measure of retinal neurodegeneration, in a subset of female and male individuals with NC (n=18F/9M), MCI (n=9F/8M) and AD (n= 33F/28M). **b** Quantification of brain atrophy severity score in a subset of female and male individuals with NC (n=11F/7M), MCI (n=9F/9M), and AD (n= 46F/41M). **c,d** Pearson correlation analyses between retinal thickness and retinal **(c)** Aβ_42_ burden, and **(d)** Oligo-tau in females versus males. **e,f** Spearman correlation analyses between brain atrophy and **(e)** brain Aβ plaque **(f)** NFTs. **g** Retinal thickness and brain atrophy stratified by sex and *APOEε4* genotype. **h** Spearman correlation between retinal thickness and brain atrophy **i-l** Spearman correlation analyses between retinal thickness or brain atrophy and **(i-j)** Braak stage, and **(k-l)** CDR scores. **m** Quantification of retinal vascular Aβ_40_ immunoreactive (IR) % area in a subset of female and male NC (n=6F/5M), MCI (n=3F/3M), and AD (n= 8F/7M) individuals. **n** CAA score in a subset of female and male NC (n=12F/8M), MCI (n=9F/9M), and AD (n= 46F/40M) individuals. Data are presented as individual subjects (circles) with group means ± SEMs. Statistical analyses used 2-way ANOVA with Bonferroni’s post hoc tests for 3- or more group comparisons and unpaired Student’s 2-tailed *t*-test for 2-group comparisons (in parentheses). Fold changes are indicated in black. **o** Heatmap showing Pearson correlation coefficient (*r_P_*), *P-value* (asterisks), and number of individuals between retinal vascular Aβ_40_ with retinal Aβ_42_ and p-tau[S396] burden, and thickness **p** Spearman correlation between retinal vascular Aβ_40_ IR % area and MMSE score. **q,r** Spearman correlation between CAA score and **(q)** brain NFTs, and **(r)** ABC score. **s** Heatmap showing Pearson’s correlation (*r_P_*), *P-value* (asterisks), and number of individuals of retinal claudin-5 (Capillary) and ZO-1 (Capillary) IR % areas with retinal Aβ_42_ burden, p-tau[S396], Oligo-tau (T22) and thickness. **t** Heatmap showing Speaman’s correlation coefficient (*r_S_*), *P-value* (asterisks), and number of individuals of retinal claudin-5 (Capillary) and ZO-1 (Capillary) IR % areas with brain Aβ plaque and NFTs severity score, Braak stage, ABC score, and MMSE score. **u,v** Spearman correlation between retinal **(u)** ZO-1 (Capillary) IR % areas and CDR score, **(v)** claudin-5 (Capillary) IR % areas and CAA score. For each correlation distribution graph, Pearson or Spearman correlation coefficient (*r*), Holm-Šídák adjusted p-values (asterisks), and number of individuals (n) are shown in the upper right corner of each graph. Black lines represent linear regression fits with 95% confidence interval. *\*P ≤ 0.05, **P ≤ 0.01, ***P ≤ 0.001, ****P ≤ 0.0001, ns: nonsignificant*.

Since AD-related neurodegeneration is closely linked to alterations in the neurovascular microenvironment^18,22,36,37,59,60^, we next examined whether the sex-specific neurodegenerative trajectories were accompanied by distinct patterns of retinal vasculopathy. We analyzed retinal vascular amyloid burden (11A50-B10^+^ vascular Aβ_40_ IR area) in sex-stratified NC, MCI, and AD donors. Retinal vascular Aβ_40_ burden increased progressively from NC to MCI and further to AD in both sexes, with substantial elevations of 8-fold in AD males (*P < 0.0001*) and 4.1-fold in AD females (*P = 0.006*) relative to NC controls (Fig. 2m). Notably, sex-stratified analysis revealed a divergent accumulation pattern, with AD males versus females exhibiting a predominant 1.7-fold increase in retinal vascular Aβ_40_ deposition (*P = 0.0135*; n = 8F/7M). Similarly, cerebral amyloid angiopathy (CAA) scores increased 2.2-fold in AD females, whereas AD males showed a more pronounced 5.1-fold increase relative to NC controls (Fig. 2n; *P = 0.02-0.0078*). Of note, in females but not males, retinal vascular Aβ_40_ burden showed strong associations with retinal Aβ_42_ burden (Fig. 2o; *r_P_* = 0.80; *P_adj_ = 0.20-0.0002*, n = 18F/14M) and with retinal thinning (*r_P_* = –0.66–0.18; *P_adj_ = 0.57-0.008*, n = 17F/14M), indicating tight coupling of vascular Aβ_40_ and Aβ_42_ deposition to retinal neurodegeneration only in females. Notably, in males but not females, retinal vascular Aβ_40_ burden was strongly correlated with cognitive impairment (Fig. 2p; *r_S_* = –0.84; *P = 0.001*), and CAA severity showed moderate-to-strong associations with cerebral NFTs and ABC scores (Fig. 2q,r; *r_S_* = 0.57–0.61; *P_adj_ < 0.0001*, n = 66-67F/56M); by contrast, in females, CAA significantly but weakly correlated with NFTs and ABC scores (*r_S_* = 0.32-0.34; *P_adj_ = 0.009*).

We previously reported significant reductions in retinal endothelial tight-junction integrity proteins—claudin-5 and zona occludens-1 (ZO-1)—in postmortem retinas from patients with MCI and AD relative to NC controls^36^. However, how these tight-junction proteins changes correlate with canonical AD pathologies and clinical trajectories in females and males separately remains unknown. Sex-stratified analysis revealed progressive loss of retinal claudin-5⁺ and ZO1^+^ tight junction proteins in both sexes; however, this reduction reached significance at the AD dementia stage only in males, not in females (Extended data Fig. 4k,l). Our sex-specific assessments further revealed that, in males but not females, retinal claudin-5 and ZO1 tight-junction proteins were more susceptible to increasing retinal Aβ_42_, pS396-tau, and T22^+^ oligomeric tau burden, and showed moderate correlations with retinal thinning (Fig. 2s; *r_P_* = 0.48-0.75; *P_adj_ < 0.05-0.001*, n = 22-25F/20-23M). An exception was the stronger association between retinal claudin-5⁺ loss and T22^+^ oligomeric tau in females than in males (*r_P_* = –0.63 and *P_adj_ = 0.002* in females vs. –0.49 and *P_adj_ < 0.04* in males). When related to cerebral pathology, significant associations were restricted to males, in whom retinal claudin-5⁺ levels, but not ZO1^+^, correlated inversely with cerebral NFT burden, Braak stage and ABC score (Fig. 2t; *r_S_* = –0.61 to –0.70; *P_adj_ = 0.02-0.008*, n = 17F/22M), whereas no such correlations were detected in females. Retinal ZO1^+^ tight-junctions integrity correlated moderately to strongly with cognitive status (MMSE, CDR) in both sexes (Fig. 2t,u; *r_S_* = –0.68–0.63; *P_adj_ = 0.02-0.009*, n = 18-20F/16-20M), with stronger effects in males. By contrast, retinal claudin-5 correlated with MMSE (but not CDR) only in females (Fig. 2t and Extended Data 4m and Extended Data 4n; *r_S_* = 0.58; *P_adj_ = 0.02*, n = 18), but not in males. Finally, CAA severity was associated with retinal claudin-5, but not ZO-1, deficiency, with a stronger association in males than females (Fig. 2v; *r* = –0.77 males and *r* = –0.58 females; *P_adj_ = 0.004–0.0002,* n = 22F/18M).

Together, these data support greater retinal neuronal vulnerability to proteinopathies (Aβ_42_, vascular Aβ_40_, p-tau and oligomeric tau) in females, particularly at more advanced AD stages, whereas retinal vascular dysfunction, including endothelial tight-junction deficiency, is more severe and tightly coupled to CAA and cognitive decline in males.

### Female-biased retinal gliosis in AD marks more severe pathology, advanced stage, and cognitive dysfunction

Cerebral inflammation plays a central role in the onset and progression of AD with marked sex dimorphism^61^. The retina parallels key inflammatory features of the brain, and robust retinal gliosis has been reported in MCI and AD patients by our group and others^23,28,31,34,39^. However, whether these neuroinflammatory responses in the retina diverge by sex has remained unclear. To address this, we analyzed retinal gliosis by immunostaining for GFAP (astrocytes and activated Müller glia) and IBA1 (microglia) in age-matched female and male donors with NC (n =16F/10M), MCI (n = 8F/9M), and AD (n = 21F/18M). Quantitative analysis demonstrated a progressive increase in retinal GFAP^+^ macrogliosis/astrogliosis across the AD continuum in both sexes, but with significantly divergent trajectories emerging from the MCI stage to AD dementia (Fig. 3a,b). Females with AD dementia exhibited a disproportionately amplified enhancement of GFAP^+^ gliosis, with a highly significant 1.7-fold increase relative to MCI females (Fig. 3b; *P < 0.0001*). In contrast, males showed a more gradual rise that did not reach statistical significance. Consistent with this divergence, GFAP^+^ gliosis was modestly but significantly higher in AD females than AD males (1.2-fold; *P = 0.041*). Retinal macrogliosis closely tracked local AD proteinopathy in both sexes, with stronger coupling in females (Fig. 3c,d). In females, retinal GFAP^+^ IR area correlated very strongly with retinal Aβ_42_ and oligomeric tau burden (*r_P_* = 0.84–0.87; *P_adj_ < 0.0001*, n = 20–44), whereas in males, these associations were moderate-to-strong (Fig. 3c,d, *r_P_* = 0.59-0.68; *P_adj_ < 0.023–0.0001*, n = 19-36).

**Fig. 3.**
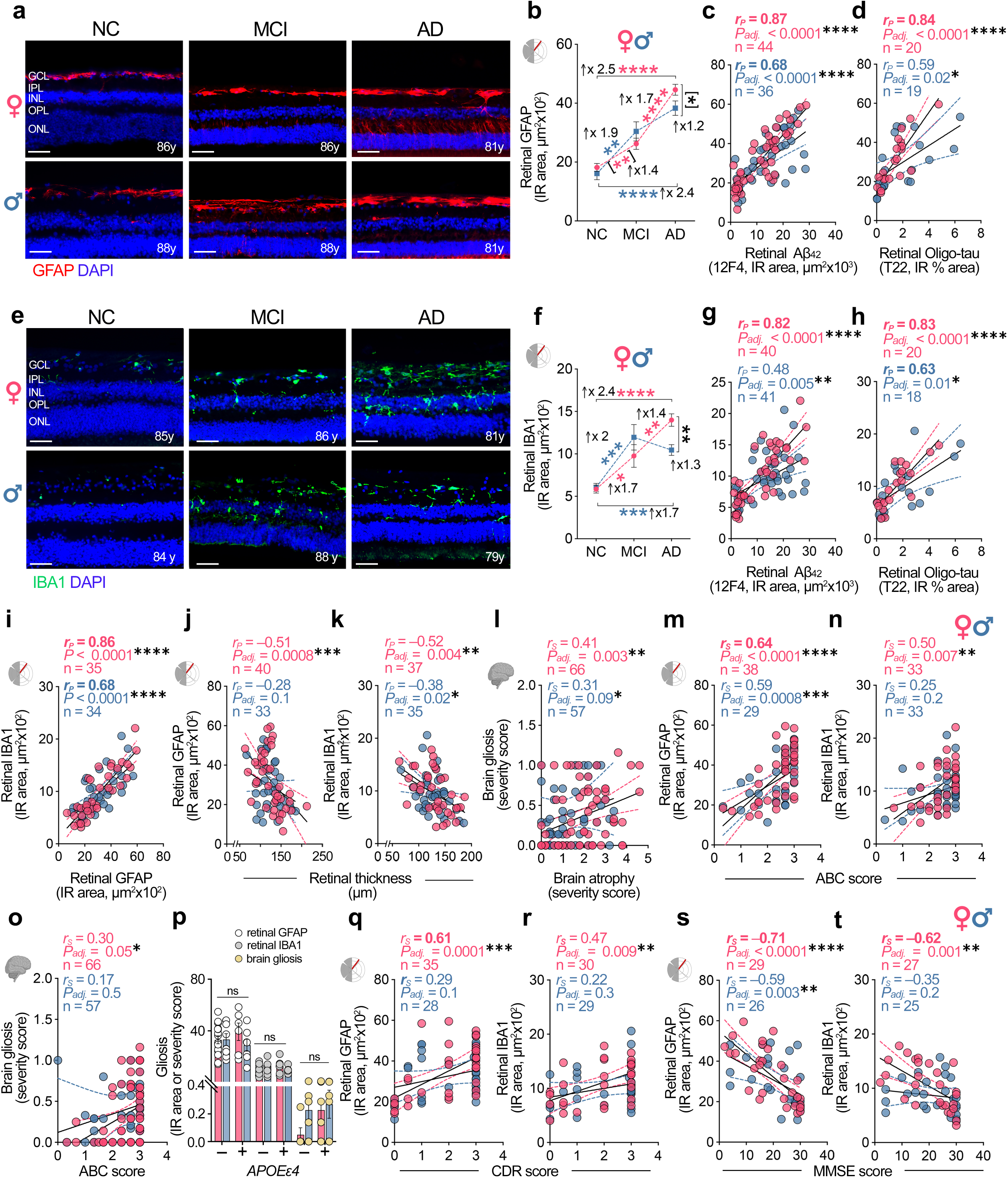
Sex-specific retinal and brain gliosis links to AD pathology and cognition. **a** Representative confocal image of postmortem retinal sections showing immunolabeling for the astrocyte and Müller glia activation marker GFAP (red, **a**), in female and male MCI, and AD patients versus NC controls. Nuclei counterstained with DAPI (blue). Scale bars: 20 µm. 4 repetitions. NFL, neurofilament layer; GCL, ganglionic cell layer; INL, inner nuclear layer; OPL, ONL, outer nuclear layer. **b** Quantification of GFAP^+^ immunoreactive (IR) areas in female and male MCI (n=8F/9M) and AD (n=21F/18M) patient retinas versus NC controls (n=16F/10M). **c,d** Pearson correlation analyses between retinal GFAP^+^ IR area and retinal **(c)** Aβ_42_, and **(d)** Oligo-tau burden. **e** Representative confocal images showing immunolabelling for microglia marker IBA1 (green), in female and male NC, MCI, and AD retinas. Nuclei counterstained with DAPI (blue). Scale bars: 20 µm. 4 repetitions. NFL, neurofilament layer; GCL, ganglionic cell layer; INL, inner nuclear layer; OPL, ONL, outer nuclear layer. **f** Quantification of IBA1^+^ IR area in female and male MCI (n=8F/9M) and AD (n=17F/23M) patient retinas versus NC controls (n=16F/10M). Fold changes are indicated in black. Data are presented as individual subjects (circles), and group means ± SEMs. 2-way ANOVA with Bonferroni’s multiple comparison tests for 3- or more group comparisons and 2-tailed unpaired Student’s *t*-test for 2-group comparisons (in parentheses). **g-i** Pearson correlation analyses between retinal IBA1^+^ IR area and retinal **(g)** Aβ_42_, **(h)** Oligo-tau burden, and **(i)** GFAP^+^ IR area. **j,k** Pearson correlation analyses between retinal GFAP^+^ or IBA1^+^ IR area and retinal thickness. **l** Spearman correlation between brain gliosis and atrophy. **m-o** Spearman correlation between retinal GFAP^+^ or IBA1^+^ IR area or brain gliosis and ABC score. **p** Retinal and brain gliosis stratified by sex and *APOEε4* genotype. **q-t** Spearman correlation between retinal GFAP^+^ or IBA1^+^ IR area and **(q,r)** CDR score, and **(s,t)** MMSE score. For each correlation analysis, Pearson or Spearman correlation coefficient (*r*), Holm-Šídák adjusted p-values (asterisks), and number of individuals (n) are shown in the upper right corner of each graph. Black lines represent linear regression fits with 95% confidence interval. *\*P ≤ 0.05, **P ≤ 0.01, ***P ≤ 0.001, ****P ≤ 0.0001*.

Retinal microgliosis, quantified as IBA1⁺ IR area, showed a similar but more pronounced sex-dependent trajectory (Fig. 3e,f). Relative to NC, males at the MCI stage exhibited a stronger early rise in retinal IBA1⁺ microgliosis (2.0-fold; *P = 0.0006*), whereas females showed a more modest but still significant increase (1.7-fold; *P = 0.0239*). Notably, this pattern diverged at AD dementia: microglial activation plateaued in males, while females demonstrated a continued acceleration, with a further 1.4-fold increase in AD versus MCI (*P = 0.009*) and an overall 1.3-fold higher IBA1⁺ burden than AD males (Fig. 3f; *P = 0.0022*). In females, retinal microgliosis was tightly coupled to local AD proteinopathy and to macroglial activation. Retinal IBA1⁺ IR area correlated very strongly with retinal Aβ_42_ and oligomeric tau burden (Fig. 3g,h; *r_P_* = 0.82–0.83; *P_adj_ < 0.0001*, n = 20–40) and showed a very strong inter-relationship with retinal GFAP⁺ macrogliosis (Fig. 3i; *r_P_* = 0.86; *P < 0.0001*, n = 35). In males, these associations were weaker overall (*r_P_* = 0.48–0.68; *P < 0.0001–0.02*, n = 18–41). Together, these data indicate that retinal glial activation scales with local proteinopathy in both sexes but is more pronounced in females.

In the brain, gliosis increased across disease stages in both sexes but did not reach statistical significance, likely reflecting the limited sensitivity and cell-type specificity of routine H&E-based clinical assessments (Extended Data Fig. 5a). To evaluate clinical relevance across CNS compartments, we next tested associations between gliosis, neurodegeneration, disease stage, and cognition in a sex-stratified manner. Retinal gliosis significantly and moderately correlated with retinal thinning in females (Fig. 3j,k; *r_P_* = –0.51 to –0.52; *P_adj_ = 0.004–0.0008*, n = 37-40), but this correlation was weak and non-significant in males. Similarly, retinal and cerebral gliosis showed weak-to-moderate correlations with brain atrophy that were more significant in females (Fig. 3l and Extended Data Fig.5b,c; *r_S_* = 0.38-0.42; *P* = *0.02-0.003*, n = 33-66; additional associations with brain Aβ plaques and NFTs are shown in Extended Data Fig.5d-i), consistent with tighter coupling between glial activation, proteinopathy, and tissue loss in females. Across neuropathologic staging, retinal GFAP⁺ gliosis showed consistently strong associations with ABC score and Braak stage in both females and males (Fig. 3m and Extended Data Fig.5j; *r_S_* = 0.59-0.65; *P ≤ 0.002-0.0001*, n = 38F/29M). In contrast, associations between retinal IBA1^+^ microgliosis or brain gliosis and ABC/Braak were generally less tight than those of retinal GFAP⁺ gliosis, but were more evident in females (Fig. 3n,o and Extended Data Fig.5k,l; *r_S_* = 0.13-0.50; *P = 0.5-0.007,* n = 33-66). Stage-stratified analyses further supported progressive increases in retinal macro-and microgliosis, as well as brain gliosis, with advancing Braak and ABC stage, with the trend more apparent in females (Extended Data Fig. 5m–r).

No significant differences in retinal and cerebral gliosis were detected between *APOEɛ4* carriers and non-carriers in either sex (Fig.3p; retina: n = 6F/7M *APOEɛ4* carriers; 16F/8M non-carriers; brain: n = 6F/9M *APOEɛ4* carriers; 13F/9M non-carriers), suggesting that *APOEε4* did not account for the heightened gliosis observed in our female cohort. Finally, retinal gliosis related to cognitive status in a strongly sex-dependent manner: retinal GFAP⁺ and IBA1⁺ measures correlated robustly with impairment (CDR and MMSE) in females (Fig. 3q-t; Females: *r_S_* = 0.47 to –0.71, *P < 0.01–0.0001*, n = 27–35), whereas these relationships were largely absent in males, except for a moderate association with MMSE (*r_S_* = –0.59; *P = 0.003*, n = 26). Brain gliosis measures were generally non-significant in both sexes, reaching significance only as a weak association with CDR in males (Extended Data Fig. 6b–I; *r_S_* = 0.37; *P = 0.02*, n = 56F/46M).

Overall, these findings reveal a female-dominant retinal neuroinflammatory phenotype in AD, characterized by sustained escalation and tighter correlation of microglial and macroglial activation to local Aβ/tau burden, neurodegeneration, and cognitive decline, particularly at advanced disease stages.

### Retinal and brain ERβ deficiency with ERα elevation tracks gliosis and AD severity in females

Considering the prominent inflammatory gliosis across the AD retina–brain axis, especially in females, we next investigated the molecular pathways that might underlie these sex-dependent inflammatory signatures. We therefore conducted an endocrine-focused reanalysis of independent human mass spectrometry–based proteomic datasets from retina (n = 6 AD, n = 6 NC) and temporal cortex (n = 10 AD, n = 8 NC). Gene Ontology (GO) analysis of differentially expressed proteins revealed significant dysregulation of estrogen receptor–associated biosynthetic and signaling networks in both tissues in AD versus NC controls (Fig. 4a,b; red arrows). In the temporal cortex, altered pathways included estrogen receptor (ESR)-mediated signaling, extranuclear estrogen signaling, nuclear receptor signaling, steroid biosynthesis and metabolism, and estrogen-dependent gene expression (Fig. 4a). The retina showed a similar pattern, with significant disruption of canonical estrogen signaling, ESR-mediated signaling, nuclear receptor signaling, steroid biosynthesis and metabolism, extranuclear estrogen signaling, and estrogen biosynthetic processes (Fig. 4b). Consistently, Ingenuity Pathway Analysis (IPA) upstream regulator prediction further identified ESR1 (ER-α) and ESR2 (ER-β) among the top sex hormone–related regulators driving these shared proteomic alterations in both compartments (Fig. 4c). Notably, ER-α and ER-β was predicted to be inhibited in both cortex and retina. Collectively, these findings point to a coordinated perturbation of estrogen receptor signaling across the AD retina–brain axis. Although luteinizing hormone was likewise predicted to be inhibited in both tissues, the broad involvement of ESR pathways in nuclear and extranuclear signaling, neuroinflammatory control, and neuroprotection provided the rationale for subsequent analysis of ER abundance and localization.

**Fig. 4.**
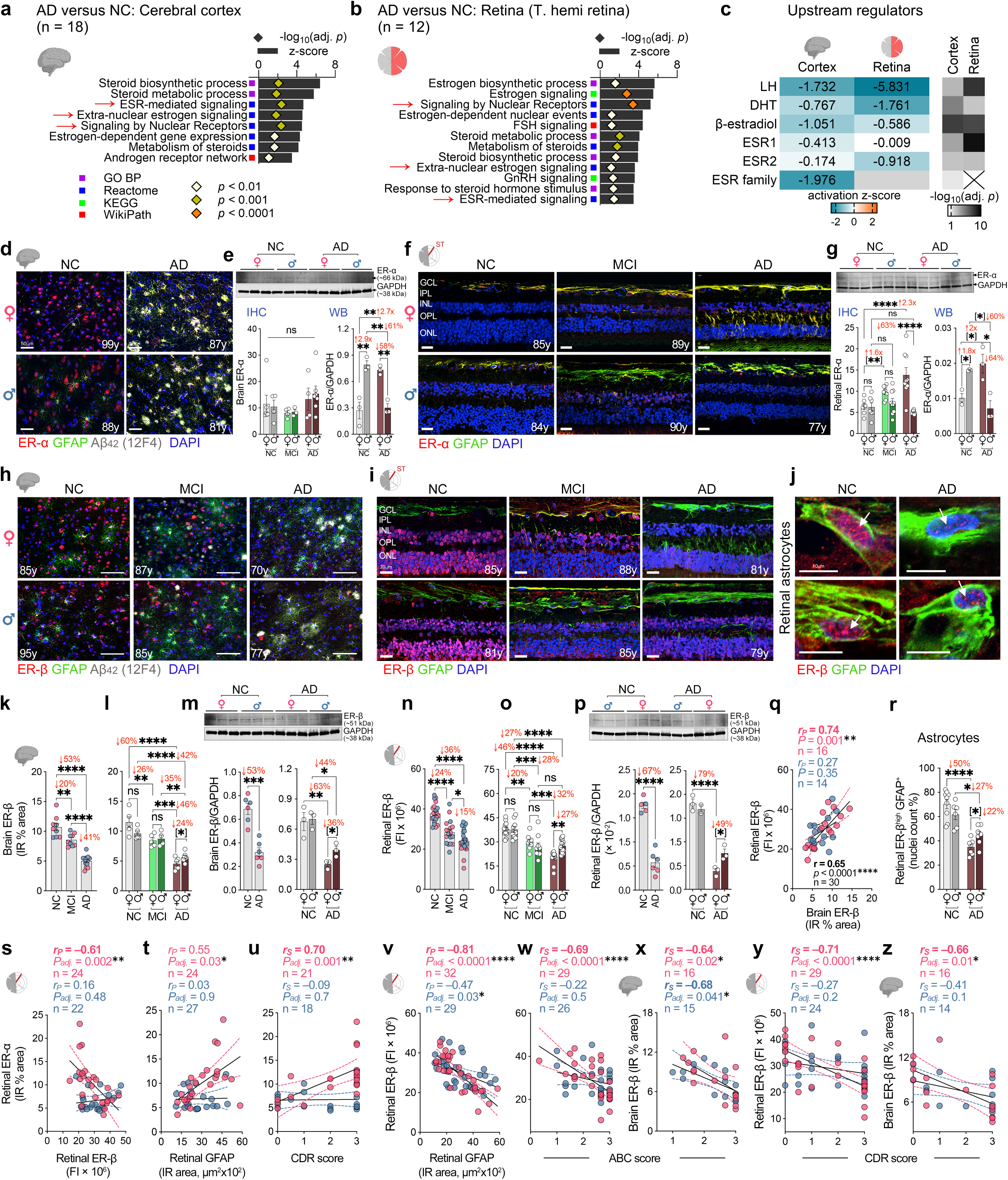
ERα and ERβ differential expression and links to disease status by sex in MCI/AD brain and retina. **a,b** Bar graphs showing dysregulated estrogen receptor-related signaling in the cerebral cortex and retina of AD patients (n=9 cortex/6 retinas) versus NC controls (n=9 cortex/6 retinas). Red arrows indicate common dysregulated pathways between cortex and retina. **c** Upstream regulators in AD cortex and retina versus NC controls. **d** Representative confocal images of postmortem cerebral cortex sections immunolabeled for ER-α (red) and astrocyte marker GFAP (green) in female and male AD patients versus NC controls. Nuclei counterstained with DAPI (blue). Scale bars: 20 µm. 2 repetitions. **e** Quantification of cortical ER-α^+^ immunoreactive (IR) % area (NC=5F/4M, MCI=4F/3M, AD=4F/6M) and WB blot analysis in female and male AD (n=3F/3M) versus NC (n=3F/3M) cortex. **f** Representative confocal images of retinal sections immunolabeled for ER-α (red) and GFAP (green) in female and male NC, MCI, and AD individuals. Nuclei counterstained with DAPI (blue). Scale bars: 20 µm. 3 repetitions. NFL, neurofilament layer; GCL, ganglionic cell layer; INL, inner nuclear layer; OPL, ONL, outer nuclear layer. **g** Quantification of retinal ER-α^+^ IR % area and WB blot analysis in female and male MCI (IF: 8F/9M) and AD (IF: 9F/8M, WB: 3F/3M) versus NC controls (IF: 8F/7M, WB: 3F/2M). **h,i** Representative confocal images of postmortem cerebral cortex and retinal sections immunolabeled for ER-β (red) and GFAP (green) in female and male MCI and AD patients versus NC controls. Nuclei counterstained with DAPI (blue). Scale bars: 20 µm. Retinal layer abbreviations are as in **f**. 2 brain and 3 retinal IF repetitions. **j** Representative confocal images showing immunolabelling for ER-β (red) in the GFAP^+^ retinal astrocytes in female and male AD and NC individuals. Nuclei counterstained with DAPI (blue). Scale bars: 10 µm, 3 repetitions. **k,l** Quantification of cortical ER-β^+^ IR % area across diagnostic groups (**k**: NC=9, MCI=9, AD=13) and by sex (**l**: NC 5F/5M, MCI 5F/4M, AD 6F/7M). **m** Western blot analysis of ER-β in AD (n=3F/3M) and NC cortex (n=3F/3M). **n,o** Quantification of retinal ER-β^+^ IR % area across diagnostic groups (**n**: NC=22, MCI=17, AD=26) and by sex (**o**: NC=13F/9M, MCI=8F/9M, AD=12F/14M). **p** Western blot analysis of retinal ER-β in AD (n=3F/3M) and NC (n=3F/2M) individuals. **q** Pearson correlation analysis between retinal and cerebral ER-β. **r** Quantification of ER-β^high^ GFAP^+^ retinal astrocytic nuclei in AD and NC individuals. **s,t** Pearson correlation analyses of retinal ER-α with retinal **(s)** ER-β and **(t)** GFAP IR % area. **u**,**v** Spearman and Pearson correlation of **(u)** retinal ER-α with CDR score, and **(v)** retinal ER-β with GFAP, respectively. **w-z** Spearman correlation analyses of retinal or cerebral ER-β with ABC and CDR scores. Fold changes are indicated in red. Data are presented as individual subjects (circles), and group means ± SEMs. 1-way ANOVA with Tukey’s or 2-way ANOVA with Bonferroni’s multiple comparison test for more than 3 groups and Student’s 2-tailed *t*-test for 2-group comparisons (in parentheses). For each correlation analysis, Pearson or Spearman correlation coefficient (*r*), Holm-Šídák adjusted *P*-values (asterisks), and number of individuals (n) are shown in the upper right corner of each graph. *\*P ≤ 0.05, **P ≤ 0.01, ***P ≤ 0.001, ****P ≤ 0.0001*.

We next determined ER-α and ER-β distribution and quantified their expression in the superotemporal (ST) retina and Brodmann area 9 (A9) cortices from NC, MCI, and AD donors (Fig. 4d–r; Extended Data Figs. 6–9). In cortex, overall ER-α levels were largely preserved across diagnostic groups, but its cellular localization was markedly altered. Compared with NC, MCI and AD cortices showed strong ER-α immunoreactivity in GFAP^+^ astrocytic processes, accompanied by weaker neuronal ER-α staining (Fig. 4d,e; Extended Data Figs. 6, 7). In NC brains, ER-α was prominent in both neuronal nuclei and cytoplasm. Notably, reactive astrocytes surrounding Aβ plaques in MCI and AD cortices exhibited intense ER-α staining, consistent with redistribution of ER-α to activated astrocytic compartments during disease progression (Extended Data Fig. 6a). Western blotting in an independent cohort further revealed pronounced sex-specific effects: cortical ER-α was increased 2.7-fold in AD females versus NC females, whereas AD males showed a 61% reduction versus NC males and 58% lower levels than AD females (Fig. 4e and Extended Data Fig. 6c; *P = 0.008-0.002*). A similar pattern emerged in retina. In MCI and AD retinas, ER-α immunoreactivity was enriched in astrocytic processes and Müller glia relative to NC, with additional labeling in the inner nuclear layer and ganglion cell layer (Fig. 4f). Retinal ER-α increased 1.6-2.3-fold in MCI and AD females relative to NC females (*P = 0.001–0.00003*), whereas AD and NC males showed comparable levels by immunostaining (Fig. 4g and Extended Data Fig. 7b). Consistent with this, total retinal ER-α by western blot did not differ significantly between AD and NC overall (Extended Data Fig. 7c), yet sex-stratified analysis revealed a 2-fold increase in AD females versus NC females, a 60% reduction in AD males versus NC males, and 64% lower ER-α in AD males than AD females (Fig. 4g; *P = 0.0496–0.011*).

In contrast to ER-α, ER-β showed a distinct trajectory across disease stages, marked by early and progressive loss in both cortex and retina (Fig. 4h–j; Extended Data Figs. 8, 9). In cortex, ER-β co-labeled with GFAP^+^ astrocytes surrounding 12F4^+^ Aβ_42_ plaques and was markedly reduced in the nuclei of both neurons and astrocytes in MCI and AD relative to NC (Fig. 4h). Quantitatively, cerebral ER-β declined by 20% in MCI and 53% in AD versus NC (Fig. 4k; *P ≤ 0.0036–0.000001*). This reduction was especially prominent in females, with cortical ER-β decreased by 26% in MCI and 60% in AD relative to NC (*P ≤ 0.0037–0.0001*), and by a further 46% in AD versus MCI (*P = 0.00012*), consistent with progressive loss across disease stages (Fig. 4l). In males, ER-β remained unchanged in MCI but declined significantly in AD, by 35% versus MCI and 42% versus NC (*P ≤ 0.003–0.00005*). Notably, cortical ER-β was 24% lower in AD females than AD males, indicating a more severe female-biased loss of ER-β signaling in AD. Western blotting confirmed these findings, showing reduced cerebral ER-β in AD versus NC overall and 36% lower ER-β in AD females than AD males (Fig. 4m; *P = 0.046*).

ER-β deficiency was also evident in the MCI and AD retina in both sexes, closely paralleling the matched brain (Fig. 4n–p). In NC retina, ER-β immunoreactivity was robust across the outer and inner nuclear layers (ONL, INL), and retinal ganglionic cell (RGC) layers (Fig. 4i). By contrast, MCI and AD retinas showed marked depletion of ER-β throughout these layers, with overall reductions of 24–36% versus NC (Fig. 4n). Sex-stratified analyses revealed retinal ER-β declines of 20–46% in females and 27–28% in males relative to their respective NC controls (Fig. 4o). In females, ER-β fell by a further 32% in AD versus MCI (*P = 0.001)*, whereas levels remained largely unchanged between MCI and AD males. As in cortex, retinal ER-β was significantly lower in AD females than AD males (27%; *P = 0.046*), indicating greater female susceptibility. Western blotting in an independent retinal cohort confirmed these findings, showing a 67% overall reduction in AD versus NC, including a striking 79% loss in AD females versus NC females and 49% lower ER-β in AD females than AD males (Fig. 4p; *P = 0.031*). Notably, retinal and cortical ER-β levels were strongly correlated in females (Fig. 4q; *r_P_* = 0.74; *P = 0.001,* n = 16F/14M), but not in males, consistent with female-biased cross-compartment ER-β vulnerability.

Co-labelling of ER-β with GFAP in the retina further revealed a pronounced depletion of nuclear ER-β within astrocytes, most prominently in females at the MCI and AD stages (Fig. 4j and 4r, and Extended data Fig. 9a). The overall proportion of ER-β⁺ astrocytes declined modestly in both sexes at the advanced AD stage (9% in females and 6% in males; *P = 0.012–0.003*) relative to their respective NC controls (Extended data Fig. 9b). However, sex-stratified analyses uncovered a substantially larger shift within the high-expressing astrocytic compartment. The proportion of retinal ER-β^high^ astrocytes declined by 50% in AD females and 27% in AD males relative to their respective NC controls (Fig. 4r; *P ≤ 0.032–0.000001*). Moreover, AD females exhibited a further 22% reduction in retinal ER-β^high^ astrocytes compared with AD males (*P = 0.045*), indicating a female-biased loss of astrocytic ER-β signaling that may contribute to the exaggerated gliosis observed in females with advanced AD.

We next examined how ER-α and ER-β in retina and brain relate to AD pathology, disease stage, *APOEε4* genotype, and cognition (Fig. 4s–z; Extended Data Figs. 10, 11). In retina, ER-β was inversely correlated with ER-α in females but not males (Fig. 4s; *r_P_* = –0.61 to 0.16; *P = 0.48–0.002*, n = 24F/22M), whereas in brain this inverse association reached significance in males (Extended Data Fig. 10t; *r_P_* = –0.57 to –0.46; *P = 0.10-0.03*, n = 13F/14M). Overall, ER-β showed the strongest and most consistent female-biased associations with clinicopathologic measures, whereas ER-α correlations were weaker and more limited (Extended Data Fig. 10 and 11). Notable exceptions were retinal ER-α, which in females correlated significantly with retinal GFAP, Braak stage, and strongly with CDR, but not in males (Fig. 4t,u; Extended Data Fig. 10n; *r* = –0.09-0.70; *P_adj_ = 0.9-0.001*, n = 14-24F/14-27M). By contrast, retinal and cerebral ER-β levels correlated moderately to very strongly with gliosis, ABC score, Braak stage, and cognitive impairment (CDR, MMSE) in females (Fig. 4v–z; Extended Data Fig. 11; *r_S_* = –0.81 to –0.64; *P_adj_ ≤ 0.02-0.0001*, n = 16-32F), whereas these relationships were overall strong to weaker or absent in males (*r_S_* = –0.68 to –0.06; *P_adj_ = 0.77-0.02*, n = 15-29M). Retinal ER-β was strongly associated with retinal Aβ_42_ and oligomeric tau consistently stronger in females (Extended Data Fig. 11a,d; *r_S_* = –0.76 to –0.71; *P_adj_ ≤ 0.002-0.0001*, n = 18-32), and its relationships with cerebral Aβ plaque and NFT burden were also significant in females (Extended Data Fig. 11b,e; *r_S_* = –0.61 to –0.51; *P_adj_ = 0.01-0.002*, n = 29) but not in males. Concordantly, cerebral ER-β deficiency linked to NFT burden in females but not males (Extended Data Fig. 11c,f; *r_S_* = –0.67 to –0.43; *P_adj_ = 0.21-0.02*, n = 16F/15M), whereas its association with cerebral Aβ plaque burden were not significant. Retinal ER-β loss also showed stronger associations in females with neurodegenerative measures, including retinal thinning and brain atrophy (Extended Data Fig. 11g-i; *r* = –0.51-0.48; *P_adj_ ≤ 0.01-0.004*, n = 29-33), whereas brain ER-β shows non-significant association. No significant differences in retinal ER-α or ER-β were detected between *APOEε4* carriers and non-carriers in either sex (Extended Data 10u and 11q), indicating that APOEε4 status does not explain the observed sex-dependent disruption of estrogen receptor signaling.

Together, these data support early and sustained disruption of estrogen receptor signaling across retina and brain in AD, in which female-biased ER-α glial reprogramming and female-selective loss of nuclear ER-β signaling may jointly amplify neuroinflammation and tighten the link between pathology and cognitive decline.

## Discussion

This comprehensive paired retina and brain study delineates sex-specific pathological trajectories across the AD continuum and determines their relationship to neuropathological and cognitive outcomes. Our integrated analyses showed substantial sex-dependent divergence in the downstream consequences of AD pathology across both retinal and cerebral compartments. Although females and males showed comparable retinal and cerebral amyloid and tau burdens, females exhibited stronger associations between retinal and cerebral proteinopathy and neurodegeneration, heightened macro- and micro-gliosis, and cognitive impairment. In contrast, males showed a dominant vasculopathic phenotype, marked by greater retinal vascular amyloid burden, accelerated cerebral amyloid angiopathy, heightened endothelial barrier susceptibility, and a stronger association with neuropathology and cognitive decline. In our cohort, we find that greater female vulnerability to AD-associated inflammation and clinical decline may stem, at least in part, from dysregulated estrogen receptor signaling across the retina and brain, with striking sex-dependent receptor remodeling. Notably, at the AD dementia stage, females showed a more pronounced increase in ER-α expression together with a marked decrease in ER-β across the retina and brain, especially in astrocytic nuclei, whereas males showed a decline in both receptors. Furthermore, ER-α and ER-β in females, but not in males, tightly correlated with glial activation and cognitive impairment. Overall, these findings reveal coordinated sex-dependent pathogenic trajectories across the retina–brain axis, implicating estrogen receptor dysregulation as molecular features of female-biased AD vulnerability, establishing the retina as a clinically accessible and mechanistically informative window into sex-specific disease biology.

Our findings suggest that sex differences in AD are less related to amyloid burden but rather to a stronger association between proteinopathy, gliosis, and cognitive decline in females. Despite similar increases in retinal and cerebral Aβ burden in females and males across diagnostic groups, with tauopathy showing only a modest tendency toward greater accumulation in females, both proteinopathies exhibited markedly stronger associations with cognitive impairment in females, with weaker or nonsignificant associations in males. This observation is consistent with recent large-scale clinical evidence from the ADNI and Mayo Clinic cohorts, showing that women exhibit greater cerebral tau load, accelerated tau propagation, and steeper cognitive decline than men despite comparable amyloid burden^62–66^. These findings are especially apparent if participants are Aβ-positive and/or *APOE*-ε4 carriers^62,67^. Together, these results suggest that females are more vulnerable to the downstream neurotoxic and inflammatory consequences of comparable AD proteinopathy.

The heightened pathological vulnerability in females was accompanied by more pronounced neurodegeneration across both retinal and cerebral compartments. Our findings align with emerging evidence indicating sex-divergent neurodegenerative patterns across the AD continuum, where women experience faster atrophy rates, greater cortical atrophy^63,68,69^, and earlier hippocampal volume loss^70–72^ after accounting for age and clinical severity. In agreement with these reports, females with AD dementia in our cohort exhibited significantly more severe retinal thinning and cerebral atrophy than males. These neurodegenerative changes were mostly linked to local Aβ but not tau burden, disease stage, and cognitive decline in females but not in males, suggesting greater neuronal susceptibility to amyloidosis in females. In contrast, males appear to follow a distinct vasculopathy-dominant trajectory, characterized by greater retinal vascular Aβ_40_ burden, accelerated progression of cerebral amyloid angiopathy, and a slightly upward trend of retinal endothelial tight-junction protein loss. This pattern is consistent with prior evidence that men with AD can exhibit higher overall CAA severity and microbleed burden^73,74^.

Notably, the strong associations between CAA or retinal vascular Aβ_40_ and endothelial barrier disruption with brain NFT burden and cognitive status in males suggest that vascular pathology may be a central contributor to cognitive impairment, particularly in men. Although the mechanisms underlying this male-biased vascular vulnerability remain unclear, mosaic loss of chromosome Y (mLOY) in hematopoietic cells, an age-related somatic alteration linked to increased AD risk^75^, may represent a plausible male-specific factor and warrants further investigation. These findings extend earlier reports of retinal vascular amyloidosis and barrier dysfunction in AD^36^ by suggesting that vascular abnormalities may be particularly informative for capturing male-specific disease vulnerability.

Our study reveals a profound sex-divergent retinal glial activation in which females display a disproportionately amplified neuroinflammatory response compared to males. While previous studies identified higher cerebral and retinal microgliosis in female AD patients^31,76^, the present data substantially extend those observations by demonstrating that retinal gliosis not only intensifies across the disease continuum but also follows distinct sex-specific trajectories. Males exhibited a trend of early rise in glial activation at MCI stage followed by a relative plateau, whereas females demonstrated a sustained escalation from MCI to AD, accompanied by stronger macroglia–microglia coupling, and markedly tighter associations with local Aβ and tau burden, retinal thinning, brain atrophy, disease stage, and cognitive impairment. Although cerebral gliosis in our cohort was assessed using non-specific H&E staining and did not differ significantly by sex, prior studies have reported higher cerebral inflammation^77,78^ and a greater burden of white-matter abnormalities in women than in men^79,80^. Together, these findings suggest that comparable proteinopathy burden triggers more severe inflammatory and neurodegenerative consequences in females, potentially mediated by female-enriched disease-associated glial states and stronger microglia-mediated interaction between amyloid and tau pathology in women^28,31,42,81,82^. These results warrant future investigation of glial cell-specific transcriptomic and proteomic signatures in the AD retina.

Importantly, our study advances an endocrine-centered mechanistic insight of sex-dependent patterns by implicating disruptions in sex hormone pathways, most notably dysregulated estrogen receptor signaling, across the retina–brain axis. Proteomic analysis identified a coordinated dysregulation of estrogen-related pathways in both the retina and cerebral cortex, and tissue-level validation revealed receptor-specific, cell-type-dependent remodeling of ER-α and ER-β expression. Among the two receptor subtypes, ER-β showed markedly greater disease-associated vulnerability than ER-α, characterized by a progressive decline across disease stages in both retina and cortex, with earlier disruption during the transition from MCI to AD and substantially greater depletion in female compared to male AD patients. This reduction was particularly prominent in females within astrocytic nuclear compartments, where ER-β loss closely paralleled stronger macroglial activation and exhibited tighter associations with amyloid burden, tau pathology, disease stage, and cognitive impairment. Notably, the ER-β deficiency was accompanied by a distinct cell-type-specific pattern for ER-α in our patient cohort. While neuronal ER-α was reduced, likely compromising direct estrogen-mediated neuroprotection and synaptic plasticity, astrocytic ER-α was markedly upregulated in both the retina and brain. This astrocytic ER-α escalation, together with progressive ER-β depletion, suggests a marked shift in estrogen-responsive signaling during disease progression. In astrocytes, increased ER-α may reflect a compensatory reactive response to mounting proteinopathy and declining ERβ-mediated anti-inflammatory control. Overall, these receptor-specific changes indicate that AD is associated not with generalized failure of estrogen signaling, but with functional redistribution of receptors that may favor reactive glial remodeling over homeostatic maintenance. This pattern was particularly pronounced in females and appeared further amplified in *APOEε4* carriers, consistent with evidence that *APOEε4* disproportionately exacerbates inflammatory vulnerability in women^83–86^. These findings may help explain why females often exhibit greater clinical vulnerability relative to males despite comparable levels of AD proteinopathy and underscore the urgent need for sex-informed therapeutic strategies.

These observations are important in the context of female aging, as decades of research suggest that the marked postmenopausal drop in estrogen and estrogen receptor signaling contributes to increased susceptibility to neuroinflammation, neurodegeneration, and faster cognitive decline in women^87^. At the same time, conventional hormone replacement therapy (HRT) using conjugated equine estrogens or estradiol has not consistently demonstrated robust neuroprotective benefit in large-scale clinical studies^88,89^. Although the ’critical window hypothesis’ proposes greater efficacy when treatment is initiated close to menopause rather than later in life, the effects on cognition remain modest and regionally variable. In this context, the observation that ER-β is depleted to a similar extent in both the retina and cortex, predominantly within astrocytic compartments, suggests that receptor-specific vulnerability, rather than global estrogen deficiency alone, may be a key determinant of heightened AD susceptibility in females. This is particularly relevant, given the established role of ER-β in restraining inflammatory responses through ligand-dependent nuclear translocation, which represses inflammatory gene transcription and promotes neuroprotective and synaptic-related gene expression, thereby supporting glial homeostasis and synaptic integrity^52,90–93^. Its preferential depletion may therefore weaken an important protective mechanism that normally limits the maladaptive neuroimmune activation, contributing to the amplified inflammatory response observed in females. The parallel loss of ER-β across both CNS compartments may also help explain why generalized estrogen replacement has yielded limited benefit. Hence, these findings support further investigation of receptor-selective strategies, particularly ER-β targeted approaches, as a potentially more mechanistically focused avenue for addressing female vulnerability in AD.

These findings also have significant translational implications. The strong concordance between retinal and cortical alterations indicates that the retina effectively mirrors core sex-dependent disease mechanisms occurring in the brain, including neuroinflammatory remodeling and selective ER-β loss. This is especially relevant because the retina is uniquely accessible in living patients, supporting its potential utility as a clinically informative site for monitoring sex-specific disease progression and stratifying disease vulnerability. The absence of a significant *APOEε4* effect on retinal ER-α or ER-β expression in our cohort further suggests that estrogen receptor dysregulation may represent a disease-related axis not solely determined by *APOEε4* genotype. Furthermore, although ER-α confers neuroprotection, its therapeutic targeting is constrained by its pro-proliferative effects in reproductive tissues and the associated risk of malignancy^52,94,95^. These limitations provide a compelling rationale for selectively targeting ER-β as a potentially safer neuroprotective strategy for women.

Several limitations of this study warrant consideration. First, as a cross-sectional postmortem investigation, the reported associations do not establish the sequence of progressive changes or direct causality. Although the observed relationships between estrogen receptor expression, glial activation, vascular compromise, and cognitive decline were robust across both retina and brain, longitudinal clinical studies and mechanistic preclinical models will be required to define the causal hierarchy through which ERα/ERβ imbalance may contribute to sex-biased disease progression. Second, although the use of paired human retinal and brain tissues provides a unique opportunity to evaluate coordinated retina–brain pathology, postmortem analyses cannot fully capture dynamic disease processes occurring in living patients. Nonetheless, by integrating paired retina and brain measures across the full clinicopathological spectrum, our study indicates that sex-dependent molecular and cellular remodeling is a fundamental, conserved feature of AD across the retina–brain axis.

In conclusion, our findings demonstrate that the retina mirrors not only core AD-related brain pathology but also sex-specific disease signatures, supporting its potential as an accessible window into sex-informed AD mechanisms. Our results indicate that amyloid and tau accumulation alone do not explain sex differences in AD, instead, comparable proteinopathy is associated with divergent downstream consequences. Females appear to be more susceptible to the neuroinflammatory and neurodegenerative consequences of AD pathology, whereas males show greater vascular injury. The close concordance between retina and brain across these processes positions the retina as a valuable site for investigating sex-specific disease biology and for developing clinically accessible biomarkers of disease severity and progression. Mechanistically, the selective depletion of ER-β across retina and brain, particularly in astrocytes, identifies ERα/ERβ imbalance a potentially important sex-linked molecular mechanism through which female vulnerability may be amplified during AD progression. These findings support further evaluation of ER-β-related signaling as a candidate therapeutic target for restoring CNS resilience during aging and AD progression.

## Methods

### Human eye and brain samples

Postmortem human eye globes and brain tissues were obtained from the Alzheimer’s Disease Research Center (ADRC) Neuropathology Core in the Department of Pathology at the University of Southern California (USC, Los Angeles, CA; IRB protocol HS-042071). In addition, eye globes were obtained from the National Disease Research Interchange (NDRI, Philadelphia, PA; IRB exempt protocol EX-1055). For a subset of patients and controls, brain specimens were also obtained from the ADRC Neuropathology Core at the University of California, Irvine (UCI [IRB protocol HS#2014–1526]). Tissue collection protocols at USC-ADRC, NDRI, and UCI-ADRC were approved by their respective oversight committees and conducted in accordance with National Institutes of Health and institutional guidelines. All the histological procedures were performed at Cedars-Sinai Medical Center under IRB protocols (Pro00053412 and Pro00019393, Pro00055802). For histological analyses, 152 retinas were collected from deceased donors with confirmed AD (n = 47F/45M) or MCI due to AD (n = 9F/9M), and from age- and sex-matched deceased donors with NC (n = 24F/18M). For protein analysis by mass spectrometry (MS), and western blot (WB), fresh-frozen cerebral cortex and retinas were obtained from an independent cohort of deceased donors comprising clinically and neuropathologically confirmed AD cases and matched NC controls. Comprehensive cohort details are provided in Table S1-3. No significant differences were observed among groups in age, sex, or postmortem interval (PMI). To protect donor confidentiality, all tissue samples were de-identified before analysis.

### Clinical and neuropathological assessments

Detailed clinical and neuropathological assessment procedures have been described in our recent publication^31,38^. Briefly, clinical and neuropathological reports, including neurological examinations and neuropsychological and cognitive assessments, were generously provided by the ADRC system via the Unified Data Set^96^. The NDRI provided patient information, including sex, ethnicity, age at death, cause of death, medical history indicating AD, the presence or absence of dementia, and any accompanying medical conditions. Most cognitive evaluations were performed annually, typically within one year before death, and for the present study, the cognitive scores closest to death were used. Clinical status was assessed using three global measures of cognition: the Clinical Dementia Rating (CDR; 0 = normal, 0.5 = very mild impairment, 1 = mild dementia, 2 = moderate dementia, 3 = severe dementia)^97^, Montreal Cognitive Assessment (MOCA scores: ≥26 = cognitively normal or <26 = cognitively impaired^98,99^, and the Mini-Mental State Examination (MMSE scores: normal cognition = 24-30; MCI = 20-23; moderate dementia = 10-19; or severe dementia ≤ 9)^100^.

Neuropathological assessment of cerebral pathology comprised evaluation of diffuse and neuritic Aβ plaques (including immature and mature forms), along with cerebral amyloid angiopathy, neurofibrillary tangles (NFTs), neuritic threads (NTs), granulovacuolar degeneration, Lewy bodies, Hirano bodies, Pick bodies, ballooned neurons, neuronal loss, microvascular changes, and gliosis. These assessments were performed across multiple brain regions, including the hippocampus, entorhinal cortex, superior frontal gyrus, superior temporal gyrus, superior parietal lobule, primary visual cortex, and visual association area in the occipital lobe. All brain samples were collected uniformly by a neuropathologist.

Formalin-fixed, paraffin-embedded (FFPE) brain sections were used to assess the severity of amyloid plaques and NFTs using anti-Aβ monoclonal antibody (mAb) clone 4G8, anti-phospho-tau mAb clone AT8, Thioflavin-S (ThioS), and Gallyas silver staining. Two neuropathologists independently scored the burden of Aβ, NFTs, and NTs using a semiquantitative scale of 0, 1, 3, and 5 [(0 = none, 1 = sparse (0-5), 3 = moderate (6-20), 5 = abundant/frequent (21-30 or above), N/A = not applicable)]. Final scores were calculated as the average of the two independent ratings. The final diagnosis included assessment of AD neuropathologic change. The Aβ plaque scores were assigned according to a system adapted from Tal et al. (A0 = no Aβ or amyloid plaques, A1 = Thal phase 1 or 2, A2 = Thal phase 3, A3 = Thal phase 4 or 5)^101^. NFT staging was based on a modified Braak classification for silver histochemistry or phospho-tau immunohistochemistry (B0 = no NFTs, B1 = Braak stage I or II, B2 = Braak stage III or IV, B3 = Braak stage V or VI)^102^. Neuritic plaque scores were adapted from CERAD (C0 = no neuritic plaques, C1 = sparse, C2 = moderate, C3 = frequent)^103^. Additional pathological features, including neuronal loss, gliosis, granulovacuolar degeneration, Hirano bodies, Lewy bodies, Pick bodies, and ballooned neurons, were evaluated using hematoxylin and eosin staining and scored as either 0 for absent or 1 for present. Cerebral amyloid angiopathy was classified into 4 grades: Grade I, amyloid deposition surrounding normal or atrophic vascular smooth muscle cells; Grade II, replacement of the vascular media by amyloid without evidence of blood leakage; Grade III, extensive amyloid deposition associated with vessel wall fragmentation and perivascular leakage; and Grade IV, extensive amyloid deposition accompanied by fibrinoid necrosis, microaneurysms, mural thrombi, luminal inflammation, and perivascular neuritis.

### Human eye and brain cortical tissues collection and processing

Donor eyes were procured with a mean post-mortem interval (PMI) of 10 hours. Upon receipt, eyes were either preserved in Optisol-GS media (Bausch & Lomb, 50006-OPT), snap frozen and stored at -80°C, or punctured once at the limbus and fixed in 10% neutral buffered formalin or 4% paraformaldehyde (PFA) and stored at 4°C. Brain tissues (Brodmann Area 9 of the prefrontal cortex) were collected from the same donors, snap frozen, and stored at -80°C. For MS, fresh-frozen human brain (medial temporal gyrus) tissues from an independent donor cohort were also obtained. Tissue collection and processing were performed using the same standardized procedures irrespective of whether donor eyes and brain tissues were obtained from NDRI, USC-ADRC, or UCI-ADRC.

### Retinal and cortical tissues preparation and cross-sections

Fresh or fixed donor eyes preserved in Optisol-GS were dissected on ice. After removal of the anterior segment to obtain eyecups, the vitreous humor was carefully removed. Retinas were isolated, separated from the underlying choroid, and prepared as flat mounts as previously described^16,31,38^. Briefly, ∼2-mm wide retinal strips extending from the ora serrata to the optic disc were dissected from four regions: superior-temporal (ST), inferior-temporal (IT), inferior-nasal (IN), and superior-nasal (SN). ST and IT strips from fixed retinas were embedded in paraffin, rotated 90° horizontally, and re-embedded for cross-sectional analysis. These retinal strips were sectioned at 8–10 μm and mounted on microscope slides coated with 3-aminopropyltriethoxysilane (APES, Sigma A3648). Temporal hemiretinas dissected from fresh tissue were stored at −80 °C for mass spectrometry (MS) and western blotting (WB). Fresh-frozen paired brain tissues from Brodmann Area 9 of the dorsolateral prefrontal cortex were fixed in 4% PFA for 16 hours at 4°C, paraffin-embedded, sectioned at 10 μm, and mounted on APES-coated slides.

### Immunofluorescence staining

#### Tissue Preparation and Antigen Retrieval

Formalin-fixed paraffin-embedded brain and retinal cross-sections were deparaffinized in xylene (2x, 10 min each), rehydrated through a graded ethanol series (100% to 70%), and rinsed in distilled water followed by 1X PBS. Heat-induced epitope retrieval was performed by incubating sections in antigen retrieval solution (pH 6.1; DAKO, S1699) at 99 °C for 1h. Retinal and brain cross-sections were washed with 1X PBS and then treated with 70% formic acid (ACROS) for 10 minutes at room temperature (RT).

#### Immunofluorescence staining

Retinal and brain sections were blocked for 1h at RT in blocking solution (DAKO, X0909) containing 0.25% Triton X-100 (Sigma, T8787), followed by overnight incubation at 4 °C with primary antibodies (Table S4). The next day, sections were washed three times in 1X PBS and incubated with fluorophore-conjugated secondary antibodies for 1h at RT (Supplementary Table 4). Slides were mounted with Fluoromount-G™ containing DAPI (Thermo Fisher, 00-4959-52). To reduce background autofluorescence, brain sections were treated with 1X TrueBlack® (Biotium, 23007) diluted in 70% ethanol (v/v) for 40 s at RT before primary antibody incubation.

### Nissl Staining

To visualize neuronal cell bodies, FFPE retinal sections were deparaffinized, rehydrated, and incubated in 0.1% Cresyl Violet acetate (Sigma, C5042) for 5 min. Following a brief rinse in running tap water, sections were dehydrated through a graded ethanol series, cleared in xylene twice (2 min each), and mounted using a xylene-based medium (Fisher Scientific, 245-691). For quantification, 12 images/section were acquired at 20X magnification, spanning the retina from the optic disc to the ora serrata. Quantitative analysis was performed to determine the percent neuronal area within the ONL and INL using Fiji.

### Microscopy and Image Analysis

Images were acquired on a Carl Zeiss Axio Imager Z1 fluorescence microscope using ZEN 2.6 Blue Edition software (Carl Zeiss MicroImaging, Inc.) or on a Nikon AX R NSPARC confocal microscope. Multichannel acquisition was used to generate composite images across fluorophore channels. For each marker and human donor, images were captured repeatedly at identical focal planes using the same exposure and gain to ensure comparability across diagnostic groups. Images were captured at 20X, 40X, and 63X objectives as required for each analysis. To sample the full retinal strip, we consistently acquired three images from the central region, 3–4 images from the mid-peripheral (M) region, and three images from the far-peripheral (F) region, as previously described^31^. Retinal thickness (µm) was measured manually using Axiovision Rel. 4.8 software, spanning from the inner limiting membrane to the outer limiting membrane. Images were exported to NIH ImageJ2/Fiji (v2.14.0) to analyze parameters of interest. For each biomarker (e.g., GFAP, IBA1, ER-α, and ER-β), single-channel 20X images (covering ∼150×10^3^ µm^2^ area/per image) were converted to grayscale and normalized to a baseline intensity using a histogram-based threshold in ImageJ2/Fiji. This baseline-derived threshold was then applied uniformly to the corresponding channel across all subjects and diagnostic groups. Particle analysis was used to quantify total immunoreactive (IR) area and percent IR area for each biomarker. Differences in retinal ER-β immunoreactivity were additionally quantified as fluorescence intensity (FI), following a previously established protocol^104^. Briefly, ER-β positive regions of interest (ROIs) were manually selected using the ImageJ freehand tool, and FI was measured after background subtraction. FI values were averaged across central, mid-peripheral, and far-peripheral regions to determine overall signal density and are presented in arbitrary units. All image processing and analysis steps were applied consistently across samples, and statistical analyses were performed on averaged FI values (mean ± SEM). For ER-β^+^ astrocytic count, nuclei co-expressing DAPI with ER-β and surrounded by GFAP astrocytic signal were manually counted using ZEN tools in the same 20X fields used for FI quantification. Nuclei were manually classified as ER-β^high^ or ER-β^weak^ based on relative nuclear ER-β fluorescence intensity. Throughout acquisition and analysis, investigators were blinded to patient diagnosis.

### Proteome Analysis by Mass Spectrometry (MS)

Secondary analyses were performed using our previously published mass spectrometry–based proteomic datasets from postmortem retina and cerebral cortex samples^31^, to identify differentially expressed proteins (DEPs) associated with endocrine hormone signaling. Briefly, the original workflow comprised: (i) preparation of retinal and cortical homogenates, (ii) tandem mass tag (TMT) labeling, (iii) nanoflow liquid chromatography coupled to electrospray ionization tandem mass spectrometry (nanoLC–ESI–MS/MS), and (iv) database searching, peptide/protein quantification, and statistical analysis, as described previously^31^.

*Pathway enrichment and upstream regulator analyses*: Functional enrichment of DEPs was performed using Metascape (https://metascape.org/) with the following criteria: overlap ≥3, *P* < 0.01, and enrichment ≥1.5. Analyses included Gene Ontology Biological Process (GO BP), Reactome, KEGG, and WikiPathways database. Enrichment results are reported as z-scores with unadjusted *P* values and Benjamini–Hochberg adjusted *P* values to control the false discovery rate (FDR). Selected pathways are shown.

Upstream regulator analysis was performed using Ingenuity Pathway Analysis (IPA; Qiagen, https://digitalinsights.qiagen.com) to identify predicted sex hormone–related regulators that may drive shared proteomic alterations in the cerebral cortex and retina of AD patients compared to NC controls. IPA activation z-scores, *P* values, and Benjamini–Hochberg adjusted *P* values are reported.

### Western blot analysis of the cerebral cortex and the retina

Fresh-frozen postmortem temporal hemiretina and cerebral cortex tissues from human donors were homogenized in radioimmunoprecipitation assay (RIPA) buffer (0.5M Tris-HCl, pH 7.4, 1.5M NaCl, 2.5% deoxycholic acid, 10% NP-40, 10mM EDTA, Millipore; 20–188) supplemented with protease and phosphatase inhibitors. Protein concentrations were determined using a bicinchoninic acid (BCA) assay kit (Pierce). Lysates were cleared with brief centrifugation for 10 min at 15000g, normalized, and boiled at 95°C after the addition of 4X SDS loading dye. Equal amounts of protein (30 µg per sample) were resolved on a 4–20% Tris-Glycine polyacrylamide gel (Bio-Rad, catalog #4561094) and transferred to polyvinylidene difluoride (PVDF) membranes. Membranes were blocked in 2.5% bovine serum albumin in 1X TBS-T (Tris-buffered saline with 0.1% Tween-20) for 1h at RT, followed by overnight incubation with primary antibodies anti-ER-α (dilution: 1:1000; cat. no. ab108398, Abcam) and anti-ER-β (dilution: 1:500; cat. no. PA1-313, Thermo Scientific) on a rocking platform at 4 °C. After washing with 1X TBS-T, membranes were incubated with fluorescent secondary antibodies, and bands were detected using the LI-COR Odyssey imaging system. Densitometric analysis was performed using Image Studio software (LI-COR), and relative protein expression was determined by normalizing target protein signals to GAPDH.

### Statistics and reproducibility

All statistical analyses were performed using GraphPad Prism (v11.0.0; GraphPad Software). Data are presented as mean ± standard error of the mean (SEM). Comparisons among three or more groups were performed using one-way or two-way analysis of variance (ANOVA), followed by Tukey’s or Bonferroni’s post hoc test for multiple comparisons. Comparisons between two independent groups were performed using two-tailed unpaired Student’s t-tests. Associations between continuous variables were assessed using Pearson’s correlation coefficient (*rₚ*) for normally distributed data or Spearman’s rank correlation coefficient (*rₛ*) for nonparametric data. Correlation coefficients and corresponding *P* values are reported to indicate the direction, strength, and significance of the associations. For multivariable correlation analyses, *P* values were adjusted for multiple comparisons using the Holm–Šídák method and reported as adjusted *P* values (*P_adj_*). *P < 0.05* was considered statistically significant. Where applicable, fold changes (FC) and/or percentage changes were calculated to estimate the magnitude and precision of observed effects.

## Acknowledgements

This work was supported by the National Institutes of Health (NIH)/the National Institute on Aging (NIA) through the following grants: R01AG056478, R01AG055865, and AG056478-04S1 (MKH). MKH is also supported by the Goldrich and Snyder Foundations. ER has been supported by the Ray Charles Foundation. AVL was supported by NEI R01 grant EY013431. We thank Drs. Rakez Kayed, Maj-Linda B. Selenica, Daniel C. Lee and late Peter Davies for sharing various anti-tau antibodies used in our previously published work. We thank Dr. Carol A. Miller for providing a portion of the human tissues and neuropathological reports and Cedars-Sinai Biobank and Pathology Core for assistance with retinal tissue FFPE processing. Illustrations Figure 1A was created with BioRender. This article is dedicated to the memory of Dr. Salomon Moni Hamaoui and Lillian Jones Black, both of whom passed away from Alzheimer’s disease.

## Author contributions

Study conception and design: SS, MKH; data acquisition: SS, YK, JPV, AR, DTF, NS, BPG, MKH; clinical assessment, brain and retinal tissue isolation and processing, as well as histological, neuropathological, and biochemical analyses: YK, SS, JPV, DTF, AR, NS, BPG, AVL, DH, LS, MKH; mass spectrometry and data analysis: JPV, MM, YK, MKH; statistical analysis: SS, JPV, DTF, AH, NS, MKH; interpretation of data and discussion: SS, MKH; manuscript writing: original draft SS, MKH; manuscript editing: SS, MKH, YK, JPV, BPG, DTF, AVL; study supervision: MKH. All authors read and approved the final manuscript.

## Competing interests

Unrelated to this study: YK, KLB, and MKH are co-founding members of NeuroVision Imaging, Inc., Sacramento, CA, USA. All other authors declare no conflict of interest related to this work.

## Reporting summary

Further information on research design is available in the Nature Portfolio Reporting Summary linked to this article.

## Data availability

Most data generated or analyzed for this study are included in this manuscript and supplementary material. Processed proteomics data are provided in the manuscript. Mass spectrometry raw files and database search results have been deposited to the ProteomeXchange Consortium via the PRIDE repository under accession no. PXD040225. Additional data supporting the findings of this study are available from the corresponding author upon reasonable request.

## Notes

### Competing Interest Statement

The authors have declared no competing interest.

## References

1. 2025 Alzheimer’s disease facts and figures. Alzheimers & Dementia 21(2025).

2. Rajan, K.B. et al. Population estimate of people with clinical Alzheimer’s disease and mild cognitive impairment in the United States (2020-2060). Alzheimers Dement 17, 1966–1975 (2021).

3. Barnes, L.L. et al. Sex differences in the clinical manifestations of Alzheimer disease pathology. Arch Gen Psychiatry 62, 685–91 (2005).

4. Filon, J.R. et al. Gender Differences in Alzheimer Disease: Brain Atrophy, Histopathology Burden, and Cognition. J Neuropathol Exp Neurol 75, 748–754 (2016).

5. Wang, Y.T. et al. Sex-specific modulation of amyloid-beta on tau phosphorylation underlies faster tangle accumulation in females. Brain 147, 1497–1510 (2024).

6. Pan, F. et al. Sex and APOE genotype differences in amyloid deposition and cognitive performance along the Alzheimer’s Continuum. Neurobiol Aging 130, 84–92 (2023).

7. Sun, Y.Y., Wang, Z. & Huang, H.C. Roles of ApoE4 on the Pathogenesis in Alzheimer’s Disease and the Potential Therapeutic Approaches. Cell Mol Neurobiol 43, 3115–3136 (2023).

8. Altmann, A., Tian, L., Henderson, V.W., Greicius, M.D. & Alzheimer’s Disease Neuroimaging Initiative, I. Sex modifies the APOE-related risk of developing Alzheimer disease. Ann Neurol 75, 563–73 (2014).

9. Guo, L., Zhong, M.B., Zhang, L., Zhang, B. & Cai, D. Sex Differences in Alzheimer’s Disease: Insights From the Multiomics Landscape. Biol Psychiatry 91, 61–71 (2022).

10. Gaire, B.P. et al. Alzheimer’s disease pathophysiology in the Retina. Prog Retin Eye Res 101, 101273 (2024).

11. Alber, J. et al. Retina pathology as a target for biomarkers for Alzheimer’s disease: Current status, ophthalmopathological background, challenges, and future directions. Alzheimers Dement 20, 728–740 (2024).

12. Doustar, J., Torbati, T., Black, K.L., Koronyo, Y. & Koronyo-Hamaoui, M. Optical Coherence Tomography in Alzheimer’s Disease and Other Neurodegenerative Diseases. Front Neurol 8, 701 (2017).

13. Du, X., et al. Label-free hyperspectral imaging and deep-learning prediction of retinal amyloid beta-protein and phosphorylated tau. PNAS Nexus 1, pgac164 (2022).

14. Hart, N.J., Koronyo, Y., Black, K.L. & Koronyo-Hamaoui, M. Ocular indicators of Alzheimer’s: exploring disease in the retina. Acta Neuropathol 132, 767–787 (2016).

15. Kelly, L. et al. Clearance of interstitial fluid (ISF) and CSF (CLIC) group-part of Vascular Professional Interest Area (PIA), updates in 2022-2023. Cerebrovascular disease and the failure of elimination of Amyloid-beta from the brain and retina with age and Alzheimer’s disease: Opportunities for therapy. Alzheimers Dement 20, 1421–1435 (2024).

16. Koronyo, Y. et al. Retinal amyloid pathology and proof-of-concept imaging trial in Alzheimer’s disease. JCI Insight 2(2017).

17. Mirzaei, N. et al. Alzheimer’s Retinopathy: Seeing Disease in the Eyes. Front Neurosci 14, 921 (2020).

18. Shi, H. et al. Retinal Vasculopathy in Alzheimer’s Disease. Front Neurosci 15, 731614 (2021).

19. Snyder, P.J. et al. Retinal imaging in Alzheimer’s and neurodegenerative diseases. Alzheimers Dement 17, 103–111 (2021).

20. den Haan, J., et al. Amyloid-beta and phosphorylated tau in post-mortem Alzheimer’s disease retinas. Acta Neuropathol Commun 6, 147 (2018).

21. Koronyo-Hamaoui, M. et al. Identification of amyloid plaques in retinas from Alzheimer’s patients and noninvasive in vivo optical imaging of retinal plaques in a mouse model. Neuroimage 54 **Suppl 1**, S204–17 (2011).

22. Shi, H. et al. Identification of early pericyte loss and vascular amyloidosis in Alzheimer’s disease retina. Acta Neuropathol 139, 813–836 (2020).

23. Xu, Q.A. et al. Muller cell degeneration and microglial dysfunction in the Alzheimer’s retina. Acta Neuropathol Commun 10, 145 (2022).

24. Cao, K.J. et al. ARCAM-1 Facilitates Fluorescence Detection of Amyloid-Containing Deposits in the Retina. Transl Vis Sci Technol 10, 5 (2021).

25. Cao, Q. et al. Transport of beta-amyloid from brain to eye causes retinal degeneration in Alzheimer’s disease. J Exp Med 221(2024).

26. den Haan, J., Morrema, T.H.J., Rozemuller, A.J., Bouwman, F.H. & Hoozemans, J.J.M. Different curcumin forms selectively bind fibrillar amyloid beta in post mortem Alzheimer’s disease brains: Implications for in-vivo diagnostics. Acta Neuropathol Commun 6, 75 (2018).

27. Dumitrascu, O.M. et al. Retinal peri-arteriolar versus peri-venular amyloidosis, hippocampal atrophy, and cognitive impairment: exploratory trial. Acta Neuropathol Commun 12, 109 (2024).

28. Grimaldi, A. et al. Neuroinflammatory Processes, A1 Astrocyte Activation and Protein Aggregation in the Retina of Alzheimer’s Disease Patients, Possible Biomarkers for Early Diagnosis. Front Neurosci 13, 925 (2019).

29. Hart de Ruyter, F.J., et al. Phosphorylated tau in the retina correlates with tau pathology in the brain in Alzheimer’s disease and primary tauopathies. Acta Neuropathol 145, 197–218 (2023).

30. Hinton, D.R., Sadun, A.A., Blanks, J.C. & Miller, C.A. Optic-nerve degeneration in Alzheimer’s disease. N Engl J Med 315, 485–7 (1986).

31. Koronyo, Y. et al. Retinal pathological features and proteome signatures of Alzheimer’s disease. Acta Neuropathol 145, 409–438 (2023).

32. La Morgia, C. et al. Melanopsin retinal ganglion cell loss in Alzheimer disease. Ann Neurol 79, 90–109 (2016).

33. Lee, S. et al. Amyloid Beta Immunoreactivity in the Retinal Ganglion Cell Layer of the Alzheimer’s Eye. Front Neurosci 14, 758 (2020).

34. Nunez-Diaz, C. et al. The fluorescent ligand bTVBT2 reveals increased p-tau uptake by retinal microglia in Alzheimer’s disease patients and App(NL-F/NL-F) mice. Alzheimers Res Ther 16, 4 (2024).

35. Schultz, N., Byman, E., Netherlands Brain, B. & Wennstrom, M. Levels of Retinal Amyloid-beta Correlate with Levels of Retinal IAPP and Hippocampal Amyloid-beta in Neuropathologically Evaluated Individuals. J Alzheimers Dis 73, 1201–1209 (2020).

36. Shi, H. et al. Retinal arterial Abeta(40) deposition is linked with tight junction loss and cerebral amyloid angiopathy in MCI and AD patients. Alzheimers Dement 19, 5185–5197 (2023).

37. Shi, H. et al. Retinal capillary degeneration and blood-retinal barrier disruption in murine models of Alzheimer’s disease. Acta Neuropathol Commun 8, 202 (2020).

38. Shi, H. et al. Identification of retinal oligomeric, citrullinated, and other tau isoforms in early and advanced AD and relations to disease status. Acta Neuropathol 148, 3 (2024).

39. Wijesinghe, P. et al. Decoding amyloid beta clearance systems at inner blood-retina barrier using three-dimensional ex vivo retinal imaging in Alzheimer’s disease. Alzheimers Dement 21, e70592 (2025).

40. Heneka, M.T. et al. Neuroinflammation in Alzheimer disease. Nat Rev Immunol 25, 321–352 (2025).

41. Sala Frigerio, C., et al. The Major Risk Factors for Alzheimer’s Disease: Age, Sex, and Genes Modulate the Microglia Response to Abeta Plaques. Cell Rep 27, 1293–1306 e6 (2019).

42. Casaletto, K.B. et al. Sex-specific effects of microglial activation on Alzheimer’s disease proteinopathy in older adults. Brain 145, 3536–3545 (2022).

43. Congdon, E.E. Sex Differences in Autophagy Contribute to Female Vulnerability in Alzheimer’s Disease. Front Neurosci 12, 372 (2018).

44. Shang, D., Wang, L., Klionsky, D.J., Cheng, H. & Zhou, R. Sex differences in autophagy-mediated diseases: toward precision medicine. Autophagy 17, 1065–1076 (2021).

45. Brinton, R.D., Yao, J., Yin, F., Mack, W.J. & Cadenas, E. Perimenopause as a neurological transition state. Nat Rev Endocrinol 11, 393–405 (2015).

46. McCarthy, M. & Raval, A.P. The peri-menopause in a woman’s life: a systemic inflammatory phase that enables later neurodegenerative disease. J Neuroinflammation 17, 317 (2020).

47. Mosconi, L. et al. In vivo brain estrogen receptor density by neuroendocrine aging and relationships with cognition and symptomatology. Sci Rep 14, 12680 (2024).

48. Lopez-Lee, C., Torres, E.R.S., Carling, G. & Gan, L. Mechanisms of sex differences in Alzheimer’s disease. Neuron 112, 1208–1221 (2024).

49. Scassellati, C. et al. Promising Intervention Approaches to Potentially Resolve Neuroinflammation And Steroid Hormones Alterations in Alzheimer’s Disease and Its Neuropsychiatric Symptoms. Aging Dis 12, 1337–1357 (2021).

50. Ocanas, S.R. et al. Microglial senescence contributes to female-biased neuroinflammation in the aging mouse hippocampus: implications for Alzheimer’s disease. J Neuroinflammation 20, 188 (2023).

51. Tawarayama, H. et al. Estrogen, via ESR2 receptor, prevents oxidative stress-induced Muller cell death and stimulates FGF2 production independently of NRF2, attenuating retinal degeneration. Exp Eye Res 248, 110103 (2024).

52. Itoh, N. et al. Estrogen receptor beta in astrocytes modulates cognitive function in mid-age female mice. Nat Commun 14, 6044 (2023).

53. Carroll, J.C. et al. Progesterone and estrogen regulate Alzheimer-like neuropathology in female 3xTg-AD mice. J Neurosci 27, 13357–65 (2007).

54. Yue, X. et al. Brain estrogen deficiency accelerates Abeta plaque formation in an Alzheimer’s disease animal model. Proc Natl Acad Sci U S A 102, 19198–203 (2005).

55. Ventura-Clapier, R. et al. Mitochondria: a central target for sex differences in pathologies. Clin Sci (Lond) 131, 803–822 (2017).

56. Zhao, L., Mao, Z., Woody, S.K. & Brinton, R.D. Sex differences in metabolic aging of the brain: insights into female susceptibility to Alzheimer’s disease. Neurobiol Aging 42, 69–79 (2016).

57. Shinohara, M. et al. Impact of sex and APOE4 on cerebral amyloid angiopathy in Alzheimer’s disease. Acta Neuropathol 132, 225–234 (2016).

58. Lopez-Lee, C., Kodama, L. & Gan, L. Sex Differences in Neurodegeneration: The Role of the Immune System in Humans. Biol Psychiatry 91, 72–80 (2022).

59. Li, M. et al. Microvascular and cellular dysfunctions in Alzheimer’s disease: an integrative analysis perspective. Sci Rep 14, 20944 (2024).

60. Zlokovic, B.V. Neurovascular pathways to neurodegeneration in Alzheimer’s disease and other disorders. Nat Rev Neurosci 12, 723–38 (2011).

61. Guldner, I.H. et al. Ageing promotes microglial accumulation of slow-degrading synaptic proteins. Nature 650, 930–941 (2026).

62. Buckley, R.F. et al. Sex, amyloid, and APOE epsilon4 and risk of cognitive decline in preclinical Alzheimer’s disease: Findings from three well-characterized cohorts. Alzheimers Dement 14, 1193–1203 (2018).

63. Cieri, F. et al. Relationship of sex differences in cortical thickness and memory among cognitively healthy subjects and individuals with mild cognitive impairment and Alzheimer disease. Alzheimers Res Ther 14, 36 (2022).

64. Jack, C.R., Jr., et al. Prevalence of Biologically vs Clinically Defined Alzheimer Spectrum Entities Using the National Institute on Aging-Alzheimer’s Association Research Framework. JAMA Neurol 76, 1174–1183 (2019).

65. Mak, E. et al. Sex-Specific Associations of alpha-Synuclein Pathology With Tau Accumulation. JAMA Netw Open 9, e260461 (2026).

66. Ramanan, V.K. et al. Association of Apolipoprotein E varepsilon4, Educational Level, and Sex With Tau Deposition and Tau-Mediated Metabolic Dysfunction in Older Adults. JAMA Netw Open 2, e1913909 (2019).

67. Koran, M.E.I., Wagener, M., Hohman, T.J. & Alzheimer’s Neuroimaging, I. Sex differences in the association between AD biomarkers and cognitive decline. Brain Imaging Behav 11, 205–213 (2017).

68. Filiatrault, M. et al. Estrogen-related receptor gene expression associates with sex differences in cortical atrophy in isolated REM sleep behavior disorder. Nat Commun 16, 9016 (2025).

69. Hua, X. et al. Sex and age differences in atrophic rates: an ADNI study with n=1368 MRI scans. Neurobiol Aging 31, 1463–80 (2010).

70. Guo, L. et al. Causal relationships between hippocampal volumetric traits and the risk of Alzheimer’s disease: a Mendelian randomization study. Brain Commun 7, fcaf030 (2025).

71. Schuff, N. et al. MRI of hippocampal volume loss in early Alzheimer’s disease in relation to ApoE genotype and biomarkers. Brain 132, 1067–77 (2009).

72. van der Flier, W.M. & Scheltens, P. Alzheimer disease: hippocampal volume loss and Alzheimer disease progression. Nat Rev Neurol 5, 361–2 (2009).

73. Arvanitakis, Z. et al. Cerebral amyloid angiopathy pathology and cognitive domains in older persons. Ann Neurol 69, 320–7 (2011).

74. Graff-Radford, J. et al. Cerebral Amyloid Angiopathy Burden and Cerebral Microbleeds: Pathological Evidence for Distinct Phenotypes. J Alzheimers Dis 81, 113–122 (2021).

75. Dumanski, J.P. et al. Mosaic Loss of Chromosome Y in Blood Is Associated with Alzheimer Disease. Am J Hum Genet 98, 1208–1219 (2016).

76. Mathys, H. et al. Single-cell transcriptomic analysis of Alzheimer’s disease. Nature 570, 332–337 (2019).

77. Garland, E.F. et al. The microglial translocator protein (TSPO) in Alzheimer’s disease reflects a phagocytic phenotype. Acta Neuropathol 148, 62 (2024).

78. Mishra, A. et al. Dynamic Neuroimmune Profile during Mid-life Aging in the Female Brain and Implications for Alzheimer Risk. iScience 23, 101829 (2020).

79. Inguanzo, A. et al. Atrophy trajectories in Alzheimer’s disease: how sex matters. Alzheimers Res Ther 17, 79 (2025).

80. Morrison, C., Dadar, M., Collins, D.L. & Alzheimer’s Disease Neuroimaging, I. Sex differences in risk factors, burden, and outcomes of cerebrovascular disease in Alzheimer’s disease populations. Alzheimers Dement 20, 34–46 (2024).

81. Wu, D., Bi, X. & Chow, K.H. Identification of female-enriched and disease-associated microglia (FDAMic) contributes to sexual dimorphism in late-onset Alzheimer’s disease. J Neuroinflammation 21, 1 (2024).

82. Xu, J. et al. Regional protein expression in human Alzheimer’s brain correlates with disease severity. Commun Biol 2, 43 (2019).

83. Fisher, D.W., Bennett, D.A. & Dong, H. Sexual dimorphism in predisposition to Alzheimer’s disease. Neurobiol Aging 70, 308–324 (2018).

84. Mak, E., Kantarci, K. & Arvanitakis, Z. Sex differences in in vivo biomarkers of neurodegenerative dementia. Front Dement 4, 1706177 (2025).

85. Neu, S.C. et al. Apolipoprotein E Genotype and Sex Risk Factors for Alzheimer Disease: A Meta-analysis. JAMA Neurol 74, 1178–1189 (2017).

86. Zeng, Q., et al. Sex and APOE genotype specific brain regional vulnerability to Alzheimer’s Disease. Geroscience (2026).

87. Flannery, J.C., Tirrell, P.S., Baumgartner, N.E. & Daniel, J.M. Neuroestrogens, the hippocampus, and female cognitive aging. Horm Behav 170, 105710 (2025).

88. Barth, C. et al. Menopausal hormone therapy and the female brain: Leveraging neuroimaging and prescription registry data from the UK Biobank cohort. Elife 13(2025).

89. Henderson, V.W. et al. Cognitive effects of estradiol after menopause: A randomized trial of the timing hypothesis. Neurology 87, 699–708 (2016).

90. Han, X. et al. Role of estrogen receptor alpha and beta in preserving hippocampal function during aging. J Neurosci 33, 2671–83 (2013).

91. Liu, F. et al. Activation of estrogen receptor-beta regulates hippocampal synaptic plasticity and improves memory. Nat Neurosci 11, 334–43 (2008).

92. Saijo, K., Collier, J.G., Li, A.C., Katzenellenbogen, J.A. & Glass, C.K. An ADIOL-ERbeta-CtBP transrepression pathway negatively regulates microglia-mediated inflammation. Cell 145, 584–95 (2011).

93. Sarvari, M. et al. Estrogens regulate neuroinflammatory genes via estrogen receptors alpha and beta in the frontal cortex of middle-aged female rats. J Neuroinflammation 8, 82 (2011).

94. Ali, S. & Coombes, R.C. Estrogen receptor alpha in human breast cancer: occurrence and significance. J Mammary Gland Biol Neoplasia 5, 271–81 (2000).

95. Cooke, P.S. et al. Stromal estrogen receptors mediate mitogenic effects of estradiol on uterine epithelium. Proc Natl Acad Sci U S A 94, 6535–40 (1997).

96. Besser, L. et al. Version 3 of the National Alzheimer’s Coordinating Center’s Uniform Data Set. Alzheimer Dis Assoc Disord 32, 351–358 (2018).

97. Morris, J.C. The Clinical Dementia Rating (CDR): current version and scoring rules. Neurology 43, 2412–4 (1993).

98. Nasreddine, Z.S., et al. The Montreal Cognitive Assessment, MoCA: a brief screening tool for mild cognitive impairment. J Am Geriatr Soc 53, 695-9 (2005).

99. Ratcliffe, L.N. et al. Classification statistics of the Montreal Cognitive Assessment (MoCA): Are we interpreting the MoCA correctly? Clin Neuropsychol 37, 562–576 (2023).

100. Folstein, M.F., Folstein, S.E. & McHugh, P.R. "Mini-mental state". A practical method for grading the cognitive state of patients for the clinician. J Psychiatr Res 12, 189–98 (1975).

101. Thal, D.R., Rub, U., Orantes, M. & Braak, H. Phases of A beta-deposition in the human brain and its relevance for the development of AD. Neurology 58, 1791–800 (2002).

102. Braak, H., Alafuzoff, I., Arzberger, T., Kretzschmar, H. & Del Tredici, K. Staging of Alzheimer disease-associated neurofibrillary pathology using paraffin sections and immunocytochemistry. Acta Neuropathol 112, 389–404 (2006).

103. Mirra, S.S. et al. The Consortium to Establish a Registry for Alzheimer’s Disease (CERAD). Part II. Standardization of the neuropathologic assessment of Alzheimer’s disease. Neurology 41, 479–86 (1991).

104. Shahin, S. et al. AAV-CRISPR/Cas9 Gene Editing Preserves Long-Term Vision in the P23H Rat Model of Autosomal Dominant Retinitis Pigmentosa. Pharmaceutics 14(2022).

