## Supplementary figures and tables for "Sex-specific retina–brain signatures link ERα/ERβ imbalance with gliosis in Alzheimer’s disease"

**Table S1.** List of human donors with demographic and neuropathological details whose brains and retinas were used in this study.
**Table S2.** List of human donors whose postmortem retinas were used for mass spectrometry and Western blot.
**Table S3.** List of human donors whose postmortem brains were used for mass spectrometry.
**Table S4.** List of antibodies for immunohistochemical and biochemical analyses.
**Extended data Fig. 1:** Sex-specific stratification of retinal and cerebral Aβ burden across disease stage, cognition, and APOEε4 genotype.
**Extended data Fig. 2:** Sex-specific stratification of retinal and cerebral pathological tau burden across disease stage, cognition, and APOEε4 genotype.
**Extended data Fig. 3:** Sex-specific associations between retinal $A\beta_{42}$/tau burden, cerebral pathology, disease stage, and cognition.
**Extended data Fig. 4:** Sex-specific associations between neurodegeneration, cerebral pathology, disease stage, and cognition.
**Extended data Fig. 5:** Sex-specific stratification and associations of retinal and cerebral gliosis across neuropathological burden, disease stages and cognitive status.
**Extended data Fig. 6:** Expression of ER-α in the cerebral cortex of NC, MCI and AD individuals.
**Extended data Fig. 7:** Expression of ER-α in the retina of NC, MCI and AD individuals
**Extended data Fig. 8:** Expression of ER-β in the cerebral cortex of NC, MCI and AD individuals.
**Extended data Fig. 9:** Expression of ER-β in the retina of NC, MCI and AD individuals.
**Extended data Fig. 10:** Sex-specific associations of retinal and cerebral ER-α with disease stage, cognition, and *APOEε4* genotype.
**Extended data Fig. 11:** Sex-specific associations of retinal and cerebral ER-β with disease stage, cognition, and *APOEε4* genotype.

**Table S1.** List of human donors with demographic and neuropathological details whose brains and retinas were used in this study.

| Diagnosis | Age (years) | Sex | PMI (hours) | APOE | Thal-A | Braak- B | CERAD- C | Analysis |
|---|---|---|---|---|---|---|---|---|
| NC1 | 93 | F | 12.00 | e3/e2 | 3 | 2 | 3 | IHC |
| NC2 | 85 | F | 4.50 | e3/e3 | 2 | 1 | 2 | IHC |
| NC3 | 99 | F | 21.00 | e3/e3 | 1 | 2 | 1 | IHC |
| NC4 | 95 | F | 7.25 | e3/e3 | 3 | 3 | 2 | IHC |
| NC5 | 95 | F | 3.15 | n.a. | 1 | 0 | 0 | IHC |
| NC6 | 92 | F | 7.00 | n.a. | 1 | 1 | 0 | IHC |
| NC7 | 97 | F | 31.00 | n.a. | 3 | 1 | 2 | IHC |
| NC8 | 102 | F | 37.00 | n.a. | 3 | 1 | 3 | IHC |
| NC9 | 100 | F | 21.50 | e3/e4 | 2 | 2 | 3 | IHC |
| NC10 | 98 | F | 21.00 | e3/e3 | 2 | 1 | 2 | IHC |
| NC11 | 95 | F | 5.30 | e3/e3 | 0 | 2 | 0 | IHC |
| NC12 | 91 | F | 22.00 | n.a. | 2 | 2 | 0.5 | IHC |
| NC13 | 93 | F | 5.60 | n.a. | n.a. | n.a. | n.a. | IHC |
| NC14 | 58 | F | 5.20 | n.a. | n.a. | n.a. | n.a. | IHC |
| NC15 | 88 | F | 5.40 | n.a. | n.a. | n.a. | n.a. | IHC |
| NC16 | 86 | F | 5.00 | n.a. | n.a. | n.a. | n.a. | IHC |
| NC17 | 76 | F | 3.60 | n.a. | n.a. | n.a. | n.a. | IHC |
| NC18 | 75 | F | 5.20 | n.a. | n.a. | n.a. | n.a. | IHC |
| NC19 | 72 | F | 5.53 | n.a. | n.a. | n.a. | n.a. | IHC/WB |
| NC20 | 76 | F | 7.30 | n.a. | n.a. | n.a. | n.a. | IHC |
| NC21 | 77 | F | 5.70 | n.a. | n.a. | n.a. | n.a. | IHC |
| NC22 | 71 | F | 7.30 | n.a. | n.a. | n.a. | n.a. | IHC |
| NC23 | 73 | F | 12.70 | n.a. | n.a. | n.a. | n.a. | IHC |
| NC24 | 70 | F | n.a. | n.a. | n.a. | n.a. | n.a. | IHC |
| NC25 | 81 | M | 7.24 | e3/e4 | 3 | 1 | 2 | IHC/WB |
| NC26 | 95 | M | 6.50 | e3/e3 | 1 | 1 | 1 | IHC/WB |
| NC27 | 76 | M | 11.25 | e3/e3 | 2 | 0 | 2 | IHC |
| NC28 | 80 | M | n.a. | n.a. | n.a. | n.a. | n.a. | IHC |
| NC29 | 58 | M | n.a. | n.a. | n.a. | n.a. | n.a. | IHC |
| NC30 | 69 | M | 6.15 | n.a. | 0 | 0 | 1 | IHC |
| NC31 | 88 | M | 5.00 | n.a. | 2 | 1 | 2 | IHC |
| NC32 | 61 | M | 6.00 | n.a. | 0 | 0 | 0 | IHC |
| NC33 | 88 | M | 19.00 | e3/e4 | 2 | 3 | 2 | IHC |
| NC34 | 77 | M | 5.20 | n.a. | n.a. | n.a. | n.a. | IHC |
| NC35 | 84 | M | 8.30 | n.a. | n.a. | n.a. | n.a. | IHC |
| NC36 | 70 | M | 7.20 | n.a. | n.a. | n.a. | n.a. | IHC |
| NC37 | 74 | M | N/A | n.a. | n.a. | n.a. | n.a. | IHC |
| NC38 | 87 | M | 7.90 | n.a. | n.a. | n.a. | n.a. | IHC |
| NC39 | 78 | M | 6.50 | n.a. | n.a. | n.a. | n.a. | IHC |
| NC40 | 73 | M | 9.40 | n.a. | n.a. | n.a. | n.a. | IHC |
| NC41 | 75 | M | 6.30 | n.a. | n.a. | n.a. | n.a. | IHC |
| NC42 | 66 | M | na | n.a. | n.a. | n.a. | n.a. | IHC |
| MCI1 | 86 | F | 18.00 | e3/e4 | 3 | 1 | 3 | IHC |
| MCI2 | 93 | F | 7.75 | e3/e3 | 3 | 2 | 2 | IHC |

| Diagnosis | Age | Sex | PMI | APOE | Thal-A | Braak- B | CERAD- C | Analysis |
|---|---|---|---|---|---|---|---|---|
| MCI3 | 94 | F | 10.50 | e3/e3 | 2 | 1 | 2 | IHC |
| MCI4 | 89 | F | 1.50 | e3/e3 | 1 | 2 | 2 | IHC |
| MCI5 | 91 | F | 4.45 | n.a. | 2 | 2 | 2 | IHC |
| MCI6 | 98 | F | 7.50 | n.a. | 2 | 3 | 2 | IHC |
| MCI7 | 87 | F | 4.00 | e3/e3 | 3 | 3 | 3 | IHC |
| MCI8 | 80 | F | 3.75 | n.a.. | 3 | 3 | 3 | IHC |
| MCI9 | 88 | F | 18.00 | e3/e3 | 2 | 3 | 3 | IHC |
| MCI10 | 93 | M | 12.50 | e3/e2 | 2 | 0 | 2 | IHC |
| MCI11 | 97 | M | 3.50 | e3/e3 | 2 | 3 | 3 | IHC |
| MCI12 | 83 | M | 3.00 | n.a. | 2 | 3 | 2 | IHC |
| MCI13 | 88 | M | 12.00 | n.a. | 1 | 2 | 2 | IHC |
| MCI14 | 80 | M | 9.00 | e3/e3 | 3 | 3 | 2 | IHC |
| MCI15 | 85 | M | 16.50 | n.a. | 0 | 1 | 3 | IHC |
| MCI16 | 75 | M | 45.00 | n.a. | 1 | 1 | 2 | IHC |
| MCI17 | 90 | M | 28.25 | e4/e2 | 1 | 1 | 1 | IHC |
| MCI18 | 99 | M | 6.70 | e3/e3 | 3 | 2 | 3 | IHC |
| AD1 | 48 | F | 4.00 | n.a. | 3 | 3 | 3 | IHC/WB |
| AD2 | 42 | F | 30.00 | e3/e3 | 3 | 3 | 3 | IHC |
| AD3 | 51 | F | 6.00 | n.a. | 3 | 3 | 3 | IHC |
| AD4 | 90 | F | 9.00 | n.a. | 2 | 3 | 3 | IHC |
| AD5 | 100 | F | 7.00 | n.a. | 2 | 3 | 3 | IHC |
| AD6 | 87 | F | 10.00 | e3/e4 | 2 | 3 | 3 | IHC |
| AD7 | 70 | F | 6.25 | n.a. | 3 | 3 | 3 | IHC |
| AD8 | 66 | F | 17.00 | e3/e3 | 3 | 3 | 3 | IHC |
| AD9 | 99 | F | 4.50 | e3/e3 | 3 | 2 | 3 | IHC |
| AD10 | 85 | F | 8.50 | e3/e3 | 3 | 3 | 3 | IHC |
| AD11 | 81 | F | 3.00 | n.a. | 3 | 3 | 3 | IHC |
| AD12 | 92 | F | 11.25 | e3/e3 | 2 | 3 | 2 | IHC |
| AD13 | 88 | F | 5.00 | n.a. | 2 | 3 | 3 | IHC |
| AD14 | 86 | F | 6.00 | e3/e4 | 3 | 3 | 2 | IHC |
| AD15 | 93 | F | 12.00 | n.a. | 2 | 2 | 2 | IHC |
| AD16 | 94 | F | 5.50 | e3/e3 | 3 | 3 | 3 | IHC/WB |
| AD17 | 81 | F | 7.50 | e3/e3 | 3 | 3 | 3 | IHC |
| AD18 | 90 | F | 19.00 | e3/e4 | 3 | 3 | 3 | IHC |
| AD19 | 93 | F | 5.50 | n.a. | 2 | 3 | 3 | IHC |
| AD20 | 87 | F | 4.50 | e3/e4 | 3 | 3 | 3 | IHC |
| AD21 | 81 | F | 4.50 | n.a. | 3 | 3 | 3 | IHC |
| AD22 | 81 | F | 5.00 | n.a. | 3 | 3 | 3 | IHC/WB |
| AD23 | 88 | F | 5.25 | e3/e4 | 3 | 3 | 3 | IHC |
| AD24 | 71 | F | 3.50 | n.a. | 3 | 3 | 3 | IHC |
| AD25 | 76 | F | 18.25 | e3/e4 | 3 | 3 | 3 | IHC |
| AD26 | 96 | F | 9.00 | n.a. | 1 | 1 | 3 | IHC |
| AD27 | 93 | F | 9.00 | e3/e3 | 3 | 2 | 2 | IHC |
| AD28 | 91 | F | 35.25 | e3/e2 | 3 | 3 | 3 | IHC |
| AD29 | 96 | F | 52.00 | e3/e3 | 3 | 3 | 3 | IHC |
| AD30 | 88 | F | 25.00 | n.a. | 2 | 3 | 3 | IHC |

| Diagnosis | Age | Sex | PMI | APOE | Thal-A | Braak- B | CERAD- C | Analysis |
|---|---|---|---|---|---|---|---|---|
| AD31 | 93 | F | 4.00 | n.a. | 3 | 3 | 3 | IHC |
| AD32 | 80 | F | 33.50 | e4/e2 | 3 | 3 | 3 | IHC |
| AD33 | 87 | F | 12.50 | e3/e3 | 3 | 3 | 3 | IHC |
| AD34 | 87 | F | 31.00 | e3/e3 | 3 | 3 | 3 | IHC |
| AD35 | 85 | F | 32.50 | e3/e3 | 2 | 3 | 3 | IHC |
| AD36 | 100 | F | 24.00 | e3/e3 | 3 | 3 | 3 | IHC |
| AD37 | 64 | F | 3.38 | n.a. | 3 | 3 | 3 | IHC |
| AD38 | 89 | F | 9.00 | n.a. | 3 | 3 | 3 | IHC |
| AD39 | 97 | F | 9.16 | n.a. | 3 | 3 | 3 | IHC |
| AD40 | 90 | F | 5.00 | n.a. | 3 | 3 | 3 | IHC |
| AD41 | 63 | F | 6.50 | n.a. | 3 | 3 | 3 | IHC |
| AD42 | 94 | F | 5.25 | n.a. | 3 | 3 | 3 | IHC |
| AD43 | 90 | F | 3.75 | n.a. | 3 | 3 | 3 | IHC |
| AD44 | 70 | F | 5.00 | e4/e4 | 2 | 2 | 2 | IHC |
| AD45 | 65 | F | 3.00 | n.a. | 3 | 3 | 3 | IHC |
| AD46 | 85 | F | 4.00 | n.a. | 2 | 3 | 3 | IHC |
| AD47 | 74 | F | n.a. | n.a. | n.a. | n.a. | n.a. | IHC |
| AD48 | 39 | M | 8.00 | n.a. | 3 | 3 | 3 | IHC |
| AD49 | 40 | M | 7.50 | e3/e3 | 3 | 2 | 3 | IHC |
| AD50 | 90 | M | 19.00 | n.a. | 3 | 3 | 3 | IHC |
| AD51 | 88 | M | 7.50 | e3/e4 | 2 | 3 | 2 | IHC/WB |
| AD52 | 77 | M | 8.25 | e3/e4 | 3 | 3 | 3 | IHC |
| AD53 | 81 | M | 6.50 | e4/e4 | 3 | 3 | 3 | IHC |
| AD54 | 83 | M | 6.00 | n.a. | 3 | 2 | 3 | IHC |
| AD55 | 65 | M | 6.75 | e4/e4 | 3 | 3 | 3 | IHC |
| AD56 | 66 | M | 11.75 | n.a. | 3 | 3 | 3 | IHC |
| AD57 | 88 | M | 7.50 | e3/e2 | 3 | 3 | 3 | IHC |
| AD58 | 90 | M | 7.00 | e3/e4 | 3 | 2 | 3 | IHC |
| AD59 | 79 | M | 6.50 | n.a. | 3 | 3 | 3 | IHC |
| AD60 | 88 | M | 4.50 | n.a. | 2 | 2 | 3 | IHC |
| AD61 | 97 | M | 5.25 | e3/e3 | 3 | 2 | 3 | IHC |
| AD62 | 92 | M | 8.50 | e3/e3 | 3 | 3 | 3 | IHC |
| AD63 | 79 | M | 6.00 | n.a. | 2 | 3 | 2 | IHC |
| AD64 | 90 | M | 4.00 | n.a. | 3 | 3 | 2 | IHC |
| AD65 | 75 | M | 9.67 | n.a. | 3 | 3 | 3 | IHC |
| AD66 | 78 | M | 10.83 | e3/e4 | 3 | 3 | 3 | IHC |
| AD67 | 85 | M | 4.33 | n.a. | 3 | 2 | 3 | IHC |
| AD68 | 69 | M | 13.00 | e3/e4 | 3 | 3 | 3 | IHC |
| AD69 | 83 | M | 17.00 | e3/e3 | 3 | 3 | 3 | IHC |
| AD70 | 88 | M | 72.00 | e3/e3 | 3 | 3 | 3 | IHC |
| AD71 | 65 | M | 62.00 | e3/e4 | 3 | 3 | 3 | IHC |
| AD72 | 86 | M | 31.50 | e4/e4 | 3 | 3 | 3 | IHC |
| AD73 | 96 | M | 7.00 | e3/e3 | 3 | 3 | 3 | IHC |
| AD74 | 99 | M | 17.75 | n.a. | 3 | 3 | 3 | IHC |
| AD75 | 94 | M | 20.00 | e3/e3 | 1 | 2 | 1 | IHC |
| AD76 | 63 | M | 23.50 | e3/e4 | 3 | 3 | 2.5 | IHC |

| Diagnosis | Age | Sex | PMI | APOE | Thal-A | Braak- B | CERAD- C | Analysis |
|---|---|---|---|---|---|---|---|---|
| AD77 | 86 | M | 15.75 | e3/e4 | 3 | 2 | 2 | IHC |
| AD78 | 59 | M | 15.00 | n.a. | 3 | 3 | 2 | IHC |
| AD79 | 72 | M | 7.00 | e3/e4 | 3 | 1 | 3 | IHC |
| AD80 | 78 | M | 8.25 | e3/e4 | 3 | 2 | 3 | IHC |
| AD81 | 65 | M | 5.00 | n.a. | 3 | 3 | 3 | IHC |
| AD82 | 83 | M | 13.00 | n.a. | 3 | 3 | 3 | IHC |
| AD83 | 71 | M | 13.67 | n.a. | 3 | 3 | 3 | IHC |
| AD84 | 87 | M | 4.50 | e3/e4 | 3 | 3 | 2 | IHC |
| AD85 | 78 | M | 9.50 | n.a. | 3 | 3 | 3 | IHC |
| AD86 | 70 | M | 4.00 | e3/e3 | 3 | 3 | 3 | IHC |
| AD87 | 86 | M | 4.00 | n.a. | 3 | 3 | 3 | IHC |
| AD88 | 69 | M | 2.50 | n.a. | 3 | 3 | 3 | IHC |
| AD89 | 85 | M | 4.83 | n.a. | n.a. | n.a. | n.a. | IHC |
| AD90 | 79 | M | 10.50 | n.a. | n.a. | n.a. | n.a. | IHC |
| AD91 | 59 | M | 7.00 | n.a. | n.a. | n.a. | n.a. | IHC |
| AD92 | 72 | M | n.a. | n.a. | n.a. | n.a. | n.a. | IHC |

A (Thal), Aβ plaque score modified from Thal; B (Braak), NFT stage modified from Braak; C (CERAD), neuritic plaque score modified from CERAD.

*Abbreviations*: AD, Alzheimer's disease dementia; APOE, apolipoprotein; IHC, immunohistology; MCI, Mild cognitive impairment; n.a., not available NC, normal cognition; PMI, post-mortem interval; WB, Western blot.

**Table S2.** List of human donors whose postmortem retinas were used for mass spectrometry and Western blot.

| Donor | Sex | Age at Death | Thal (A) | Braak (B) | CERAD (C) | Braak Stage | APOE Status | Analysis |
|---|---|---|---|---|---|---|---|---|
| AD1 | F | 48 | 3 | 3 | 3 | 5.5 | n.a. | MS |
| AD2 | F | 93 | 2 | 3 | 3 | 5 | n.a. | MS |
| AD3 | M | 88 | 2 | 3 | 2 | 5.5 | e3/e4 | MS |
| AD4 | M | 88 | 1 | 2 | 2 | 3 | n.a. | MS |
| AD5 | F | 94 | 3 | 3 | 3 | 5.5 | e3/e3 | MS |
| AD6 | F | 100 | 2 | 3 | 3 | 5.5 | n.a. | MS |
| AD7 | F | 74 | n.a. | n.a. | n.a. | n.a. | n.a. | WB |
| NC1 | M | 81 | 3 | 1 | 2 | 1.5 | e3/e4 | MS |
| NC2 | F | 75 | n.a. | n.a. | n.a. | n.a. | n.a. | MS/WB |
| NC3 | F | 72 | n.a. | n.a. | n.a. | n.a. | n.a. | MS |
| NC4 | M | 69 | n.a. | n.a. | n.a. | n.a. | n.a. | MS/WB |
| NC5 | F | 79 | n.a. | n.a. | n.a. | n.a. | n.a. | MS/WB |
| NC6 | M | 85 | 0 | 1 | 3 | 1.5 | n.a. | MS |

A (Thal), Aβ plaque score modified from Thal; B (Braak), NFT stage modified from Braak; C (CERAD), neuritic plaque score modified from CERAD.

*Abbreviations*: AD, Alzheimer's disease dementia; APOE, apolipoprotein; IHC, immunohistology; MCI, Mild cognitive impairment; n.a., not available NC, normal cognition.

**Table S3.** List of human donors whose postmortem brains were used for mass spectrometry.

| Donor | Sex | Age at Death | Plaque Stage | Braak Stage | Neuritic Plaque | APOE Status |
|---|---|---|---|---|---|---|
| AD1 | M | 86 | C | 6 | 4 | e3/e4 |
| AD2 | F | 89 | C | 6 | 4 | e3/e4 |
| AD3 | F | 95 | C | 6 | 3 | e2/e3 |
| AD4 | F | 93 | C | 6 | 4 | e3/e3 |
| AD5 | F | 88 | C | 5 | 4 | e3/e3 |
| AD6 | M | 92 | C | 5 | 4 | e3/e3 |
| AD7 | M | 86 | C | 5 | 4 | e3/e3 |
| AD8 | F | 98 | C | 6 | 4 | e3/e3 |
| AD9 | F | 91 | C | 6 | 4 | e3/e3 |
| AD10 | F | 82 | C | 6 | 4 | e3/e3 |
| NC1 | F | 87 | 0 | 2 | 1 | e2/e3 |
| NC2 | F | 91 | 0 | 4 | 1 | e2/e3 |
| NC3 | F | 95 | 0 | 2 | 1 | e3/e3 |
| NC4 | F | 90 | B | 3 | 3 | e3/e3 |
| NC5 | M | 96 | 0 | 2 | 1 | e2/e3 |
| NC6 | M | 94 | 0 | 1 | 1 | e3/e3 |
| NC7 | M | 86 | 0 | 2 | 2 | e3/e3 |
| NC8 | F | 91 | A | 2 | 2 | e3/e3 |

Plaque Stage: 0, none; A, phase I-II; B, phase III; C, phase IV-V.

Neuritic Plaque scores: 4, frequent NP; 3, moderate NP; 2, sparce NP; 1, no NPs.

*Abbreviations*: AD, Alzheimer's disease; APOE, apolipoprotein; NC, normal cognition. The original human brain and retinal mass spectrometry data on this cohort were previously published in Koronyo et al., Acta Neuropathologica 2023[1].

**Table S4.** List of antibodies for immunohistochemical and biochemical analyses.

| Antibodies | Source species | Target species | Dilution | Application | Source | Catalog # |
|---|---|---|---|---|---|---|
| **Primary Antibodies** | | | | | | |
| GFAP pAb | Goat | Hu, ms | 1:500 | IHC-F | Invitrogen | 13-0300 |
| IBA1 mAb | Rabbit | Hu, ms | 1:400 | IHC-F | Wako | 019-19741 |
| IBA1 mAb | Goat | Hu, ms | 1:300 | IHC-F | NovusBio | NB100-1028 |
| $A\beta_{42}$ (12F4) mAb | Mouse | Hu | 1:1000 | WB | Biolegend | 805501 |
| $A\beta_{40}$ (11A50-B10) mAb | Mouse | Hu | 1:200 | IHC-F | Biolegend | 805401 |
| Tau Oligomers (T22) mAb | Rabbit | Hu | 1:200 | IHC-F | Dr. Rakez Kayed | - |
| Phospho-tau (Ser396) pAb | Rabbit | Hu | 1:500 | IHC-F | Anaspec | AS-54977 |
| MC-1 | Mouse | Hu | 1:200 | IHC-F | Dr. Peter Davies | - |
| Claudin-5 mAb | Mouse | Hu, ms | 1:20 | IHC-F | ThermoFisher | 35-2500 |
| Zonula Occluden-1 (ZO1) pAb | Rabbit | Hu, ms | 1:50 | IHC-F | ThermoFisher | 61-7300 |
| ER-α mAb | Mouse | Hu, ms | 1:100-1:1000 | IHC/WB | ThermoFisher | MA1-80216 |
| ER-β pAb | Rabbit | Hu, ms | 1:100-1:1000 | IHC/WB | ThermoFisher | PA1-313 |
| GAPDH (D16H11) mAb | Rabbit | Hu, ms | 1:1000 | WB | Cell signaling | 5174 |
| GAPDH mAb | Mouse | Hu, ms | 1:1000 | WB | Millipore Sigma | G8795 |
| **Secondary Antibodies** | | | | | | |
| Cy3 (anti-mouse, -rat, -goat, & -rabbit) | Donkey | | 1:200 | IF | Jackson ImmunoResearch Laboratories | 715-165-150, 112-165-167, 705-165-147, 711-165-152 |
| Cy5 (anti-mouse, -rat, -goat, & -rabbit) | Donkey | | 1:200 | IF | Jackson ImmunoResearch Laboratories | 715-175-150, 712-175-153, 705-175-147, 711-175-152 |
| Cy2 (anti-goat, & -rabbit) | Donkey | | 1:200 | IF | Jackson ImmunoResearch Laboratories | 705-225-147, 711-225-152 |
| IRDye® 680RD pAb | Rabbit | Hu, ms | 1:10000 | WB | Licorbio | 926-68071 |
| IRDye® 800CW pAb | Rabbit | Hu, ms | 1:10000 | WB | Licorbio | 926-32211 |
| IRDye® 680RD pAb | Mouse | Hu, ms | 1:10000 | WB | Licorbio | 926-68070 |
| IRDye® 800CW pAb | Mouse | Hu, ms | 1:10000 | WB | Licorbio | 926-32210 |

*Abbreviations*: Aβ - amyloid β-protein; GFAP - glial fibrillary acidic protein; IBA1 - ionized calcium binding adaptor molecule 1; IHC - immunohistochemistry; IHC-F - fluorescence; Hu - human; ms - mouse; mAb - monoclonal antibody; pAb - polyclonal antibody. WB - Western blot.

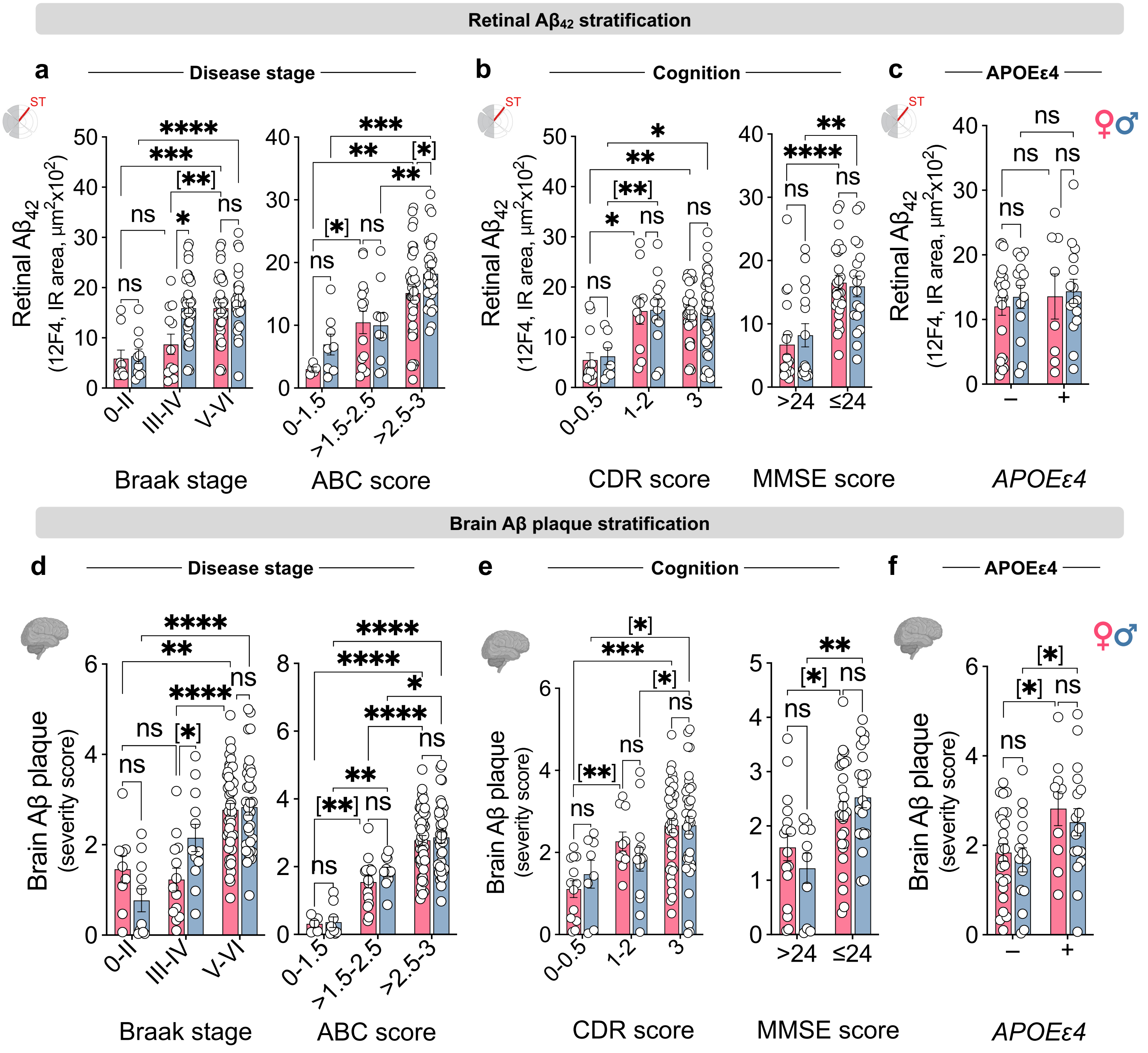


**Extended data Fig. 1: Sex-specific stratification of retinal and cerebral Aβ burden across disease stage, cognition, and *APOEε4* genotype. a-c** Retinal 12F4⁺ $A\beta_{42}$ immunoreactivity (IR) area in females and males stratified by **a** Braak stage (n = 9F/10M, 0–II; 12F/12M, III–IV; 38F/26M, V–VI) and ABC score (n = 5F/8M, 0–1.5; 15F/10M, >1.5–2.5; 39F/30M, >2.5–3) **b** cognitive status based on CDR (n = 13F/7M, 0-0.5; 13F/10M, 1-2; 31F/36M, 3) and MMSE score (n = 18F/16M, >24; 27F/17M, ≤24) and **c** *APOEε4* genotype (n = 8F/15M ε4 carriers; 24F/13M, non-carriers). **d-f** Cerebral Aβ burden in females and males stratified by **d** Braak stage (n = 9F/11M, 0–II; 13F/12M, III–IV; 45F/34M, V–VI) and ABC score (n = 5F/9M, 0–1.5; 16F/10M, >1.5–2.5; 46F/38M, >2.5–3) **e** CDR (n = 13F/8M, 0-0.5; 10F/14M, 1-2; 38F/35M, 3) and MMSE score (n = 17F/12M, >24; 27F/20M, ≤24) and **f** *APOEε4* genotype (n = 10F/17M ε4 carriers; 25F/15M, non-carriers). Data are presented as individual subjects (circles), and group means ± SEMs. Statistical analyses used 2-way ANOVA with Bonferroni's post hoc tests for 3- or more group comparisons and 2-tailed unpaired Student's *t*-test for 2-group comparisons (in parentheses). **P ≤ 0.05, **P ≤ 0.01, ***P ≤ 0.001, ****P ≤ 0.0001, ns: nonsignificant.*

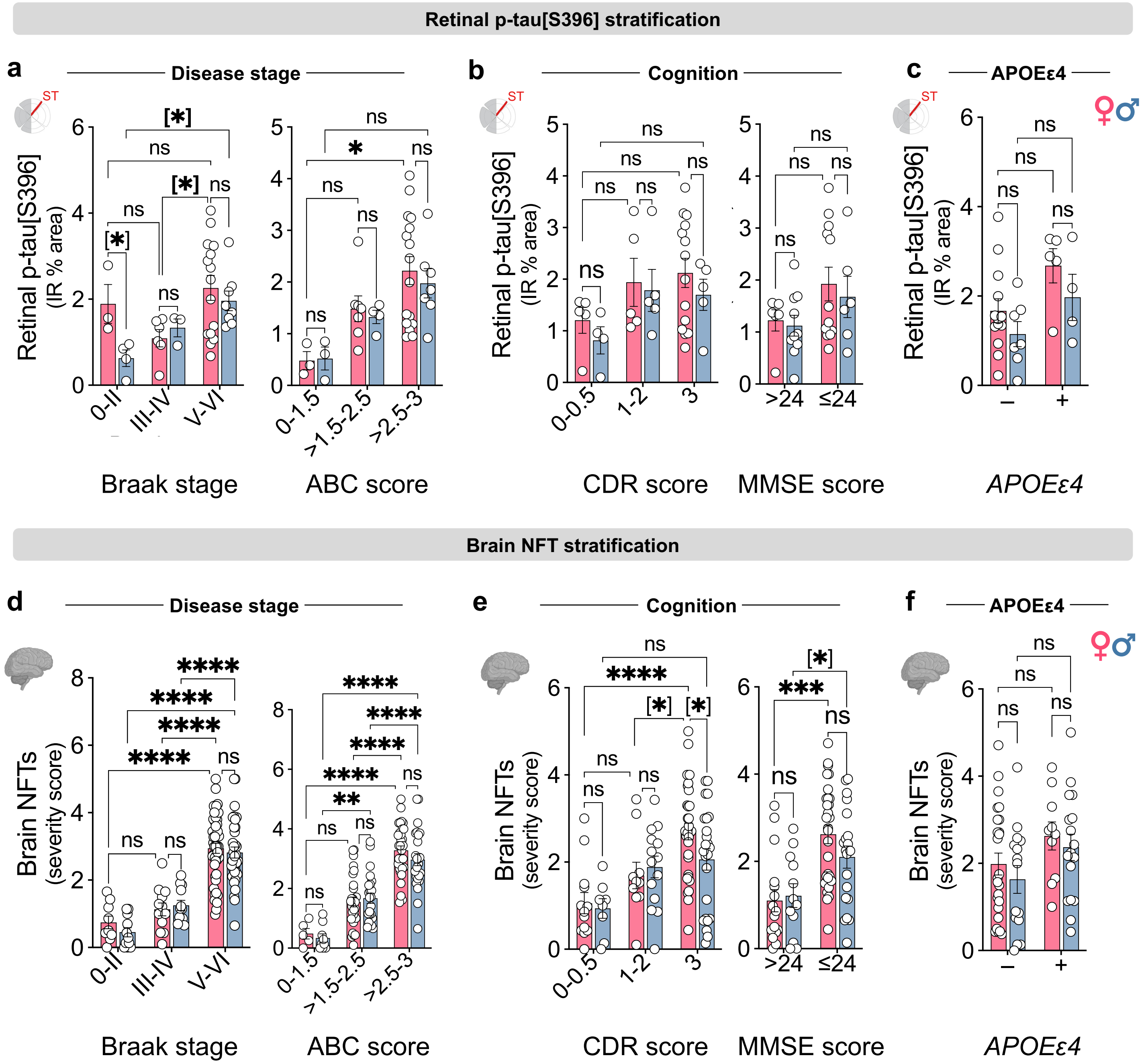


**Extended data Fig. 2: Sex-specific stratification of retinal and cerebral pathological tau burden across disease stage, cognition, and *APOEε4* genotype. a-c** Retinal p-tau[S396] immunoreactivity (IR) percentage area in females and males stratified by **a** Braak stage (n = 3F/4M, 0–II; 6F/3M, III–IV; 15F/8M, V–VI) and ABC score (n = 3F/3M, 0–1.5; 7F/4M, >1.5–2.5; 16F/7M, >2.5–3) **b** CDR (n = 5F/4M, 0-0.5; 5F/5M, 1-2; 14F/5M, 3) and MMSE scores (n = 6F/10M, >24; 12F/6M, ≤24) and **c** *APOEε4* genotype (n = 5F/4M ε4 carriers; 13F/7M, non-carriers). **d-f** Cerebral NFTs burden in females and males stratified by **d** Braak stage (n = 9F/11M, 0–II; 12F/12M, III–IV; 46F/34M, V–VI) and ABC score (n = 5F/9M, 0–1.5; 29F/23M, >1.5–2.5; 33F/25M, >2.5–3) **e** CDR (n = 13F/8M, 0-0.5; 10F/14M, 1-2; 31F/24M, 3) and MMSE score (n = 16F/12M, >24; 29F/20M, ≤24) and **f** *APOEε4* genotype (n = 10F/17M ε4 carriers; 25F/15M, non-carriers). Data are presented as individual subjects (circles), and group means ± SEMs. Statistical analyses used 2-way ANOVA with Bonferroni's post hoc tests for 3- or more group comparisons and 2-tailed unpaired Student's *t*-test for 2-group comparisons (in parentheses). **P ≤ 0.05, **P ≤ 0.01, ***P ≤ 0.001, ****P ≤ 0.0001, ns: nonsignificant.*

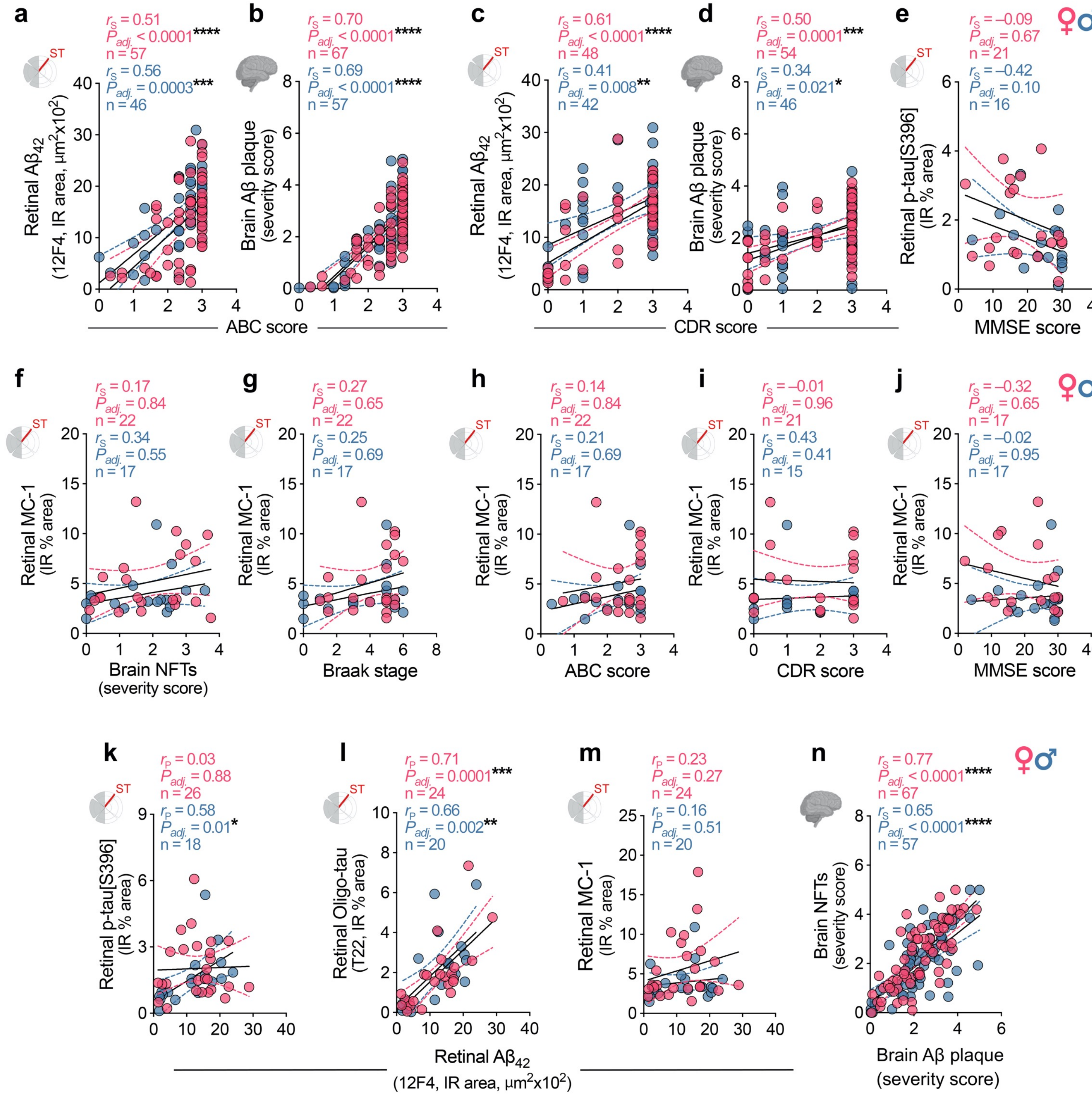


**Extended data Fig. 3: Sex-specific associations between retinal A$\beta_{42}$/tau burden, cerebral pathology, disease stage, and cognition.** **a-d** Spearman's correlation analyses between retinal or cerebral Aβ burden with **(a,b)** ABC and **(c,d)** CDR scores. **e** Spearman's correlation analysis between retinal p-tau[S396] and MMSE score. **f-j** Spearman's correlation between retinal MC1$^{+}$ pretangles and mature tangles IR % area and **(f)** brain NFTs severity score, **(g)** Braak stage, **(h)** ABC score, **(i)** CDR and **(j)** MMSE scores. **k-m** Pearson's correlation analyses between retinal A$\beta_{42}$ and retinal **(k)** p-tau[S396]**, (l)** oligomeric (Oligo)-tau and **(m)** MC1$^{+}$ pretangles and mature tangles IR % area. **n** Spearman's correlation analysis between brain NFTs severity score and brain Aβ plaque burden. Data are presented as individual values (circles). For each correlation plot, Spearman or Pearson correlation coefficient (*r*), Holm-Šídák adjusted *P* values (asterisks), and number of individuals (n) are shown in the upper right corner. Black lines represent linear regression fits with 95% confidence interval. **P ≤ 0.05, **P ≤ 0.01, ***P ≤ 0.001, ****P ≤ 0.0001, ns: nonsignificant.*

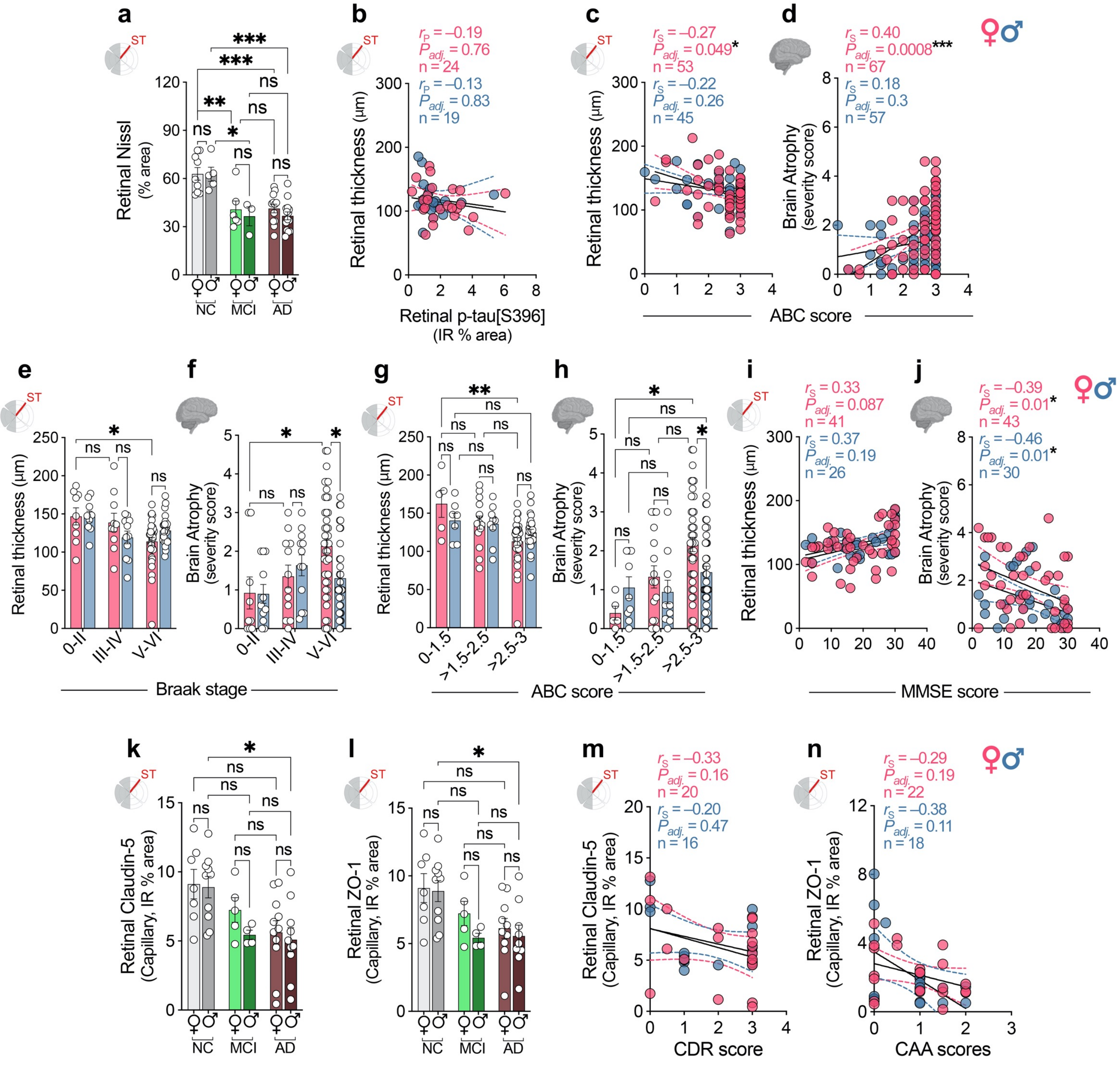

**Extended data Fig. 4: Sex-specific associations between neurodegeneration, cerebral pathology, disease stage, and cognition. a** Quantitative analysis of Nissl⁺ % area of retinas in a subset of female and male individuals with NC (n=9F/5M), MCI (n=6F/3M) and AD (n= 12F/14M). **b** Pearson's correlation between retinal thickness and p-tau[S396]. **c,d** Spearman's correlation of retinal thickness and brain atrophy with ABC score. **e-j** Retinal thickness and brain atrophy severity score stratified by **(e,f)** Braak stage (retina: n = 9F/9M, 0–II; 10F/11M, III–IV; 34F/25M, V–VI, brain: n = 9F/11M, 0–II; 12F/12M, III–IV; 46F/34M, V–VI) and **(g,h)** ABC score (retina: n = 5F/7M, 0–1.5; 14F/10M, >1.5–2.5; 34F/28M, >2.5–3 brain: n = 5F/9M, 0–1.5; 16F/10M, >1.5–2.5; 46F/38M, >2.5–3). **i,j** Spearman's correlation of retinal thickness and brain atrophy with MMSE score. **k,l** Quantitative analysis of Claudin-5 (NC = 7F/10M, MCI = 5F/4M, AD = 11F/10M) and ZO-1 (NC = 7F/10M, MCI = 5F/4M, AD = 10F/9M) IR % area in females and males across diagnostic groups. **m,n** Spearman's correlation between retinal **(m)** Claudin-5 and CDR score and **(n)** ZO-1 and CAA scores. Data are presented as individual subjects (circles), and group means ± SEMs. Statistical analyses used 2-way ANOVA with Bonferroni's post hoc tests. For each correlation plot, Pearson or Spearman correlation coefficient (*r*), Holm-Šídák adjusted *P* values (asterisks), and number of individuals (n) are shown in the upper right corner. Black lines represent linear regression fits with 95% confidence interval. **P ≤ 0.05, **P ≤ 0.01, ***P ≤ 0.001, ****P ≤ 0.0001, ns: nonsignificant.*

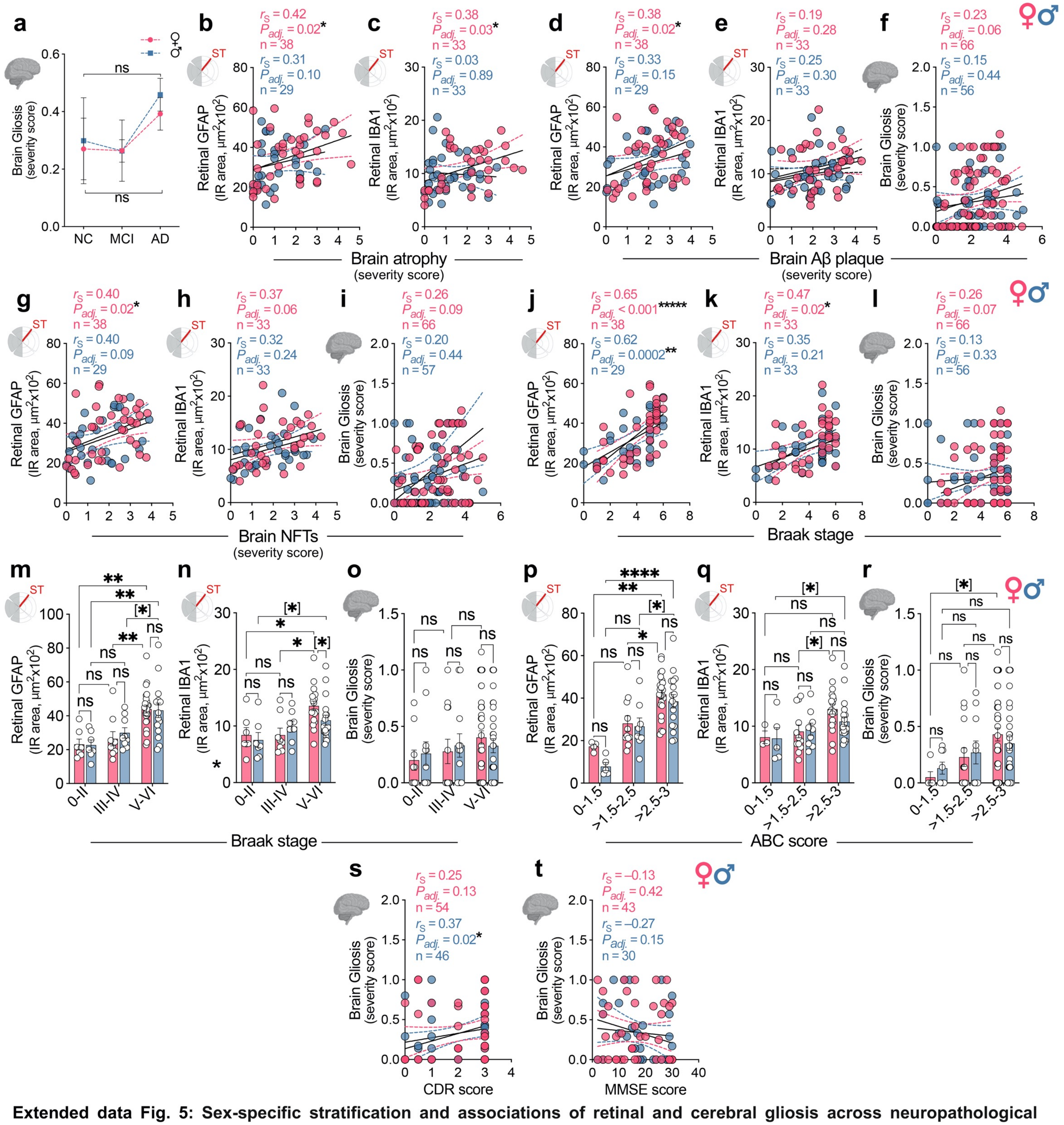


**Extended data Fig. 5: Sex-specific stratification and associations of retinal and cerebral gliosis across neuropathological burden, disease stages and cognitive status.** **a** Brain gliosis severity score in MCI, AD and NC individuals. **b,c** Spearman's correlation between retinal GPAF or IBA1 IR area with brain atrophy. **d-l** Spearman's correlation of retinal GPAF, IBA1, cerebral gliosis with **(d-f)** brain Aβ plaque **(g-i)** brain NFTs, and **(j-l)** Braak stage. **m-r** Retinal GFAP$^+$ and IBA1$^+$ immunoreactivity (IR) area, and cerebral gliosis in females and males stratified by **(m-o)** Braak stage (n = 7F/7M, 0–II; 9F/9M, III–IV; 24F/15M, V–VI) and **(p-r)** ABC score (n = 4F/5M, 0–1.5; 11F/8M, >1.5–2.5; 24F/17M, >2.5–3). **s,t** Spearman's correlation between cerebral gliosis and **(s)** CDR score, **(t)** MMSE score. Data are presented as individual subjects (circles), and group means ± SEMs. Statistical analyses used 2-way ANOVA with Bonferroni's post hoc tests for 3- or more group comparisons and 2-tailed unpaired Student's *t*-test for 2-group comparisons (in parentheses). For each correlation plot, Pearson or Spearman correlation coefficient (*r*), Holm-Šídák adjusted *P* values (asterisks), and number of individuals (n) are shown in the upper right corner. Black lines represent linear regression fits with 95% confidence interval. **P ≤ 0.05, **P ≤ 0.01, ***P ≤ 0.001, ****P ≤ 0.0001, ns: nonsignificant.*

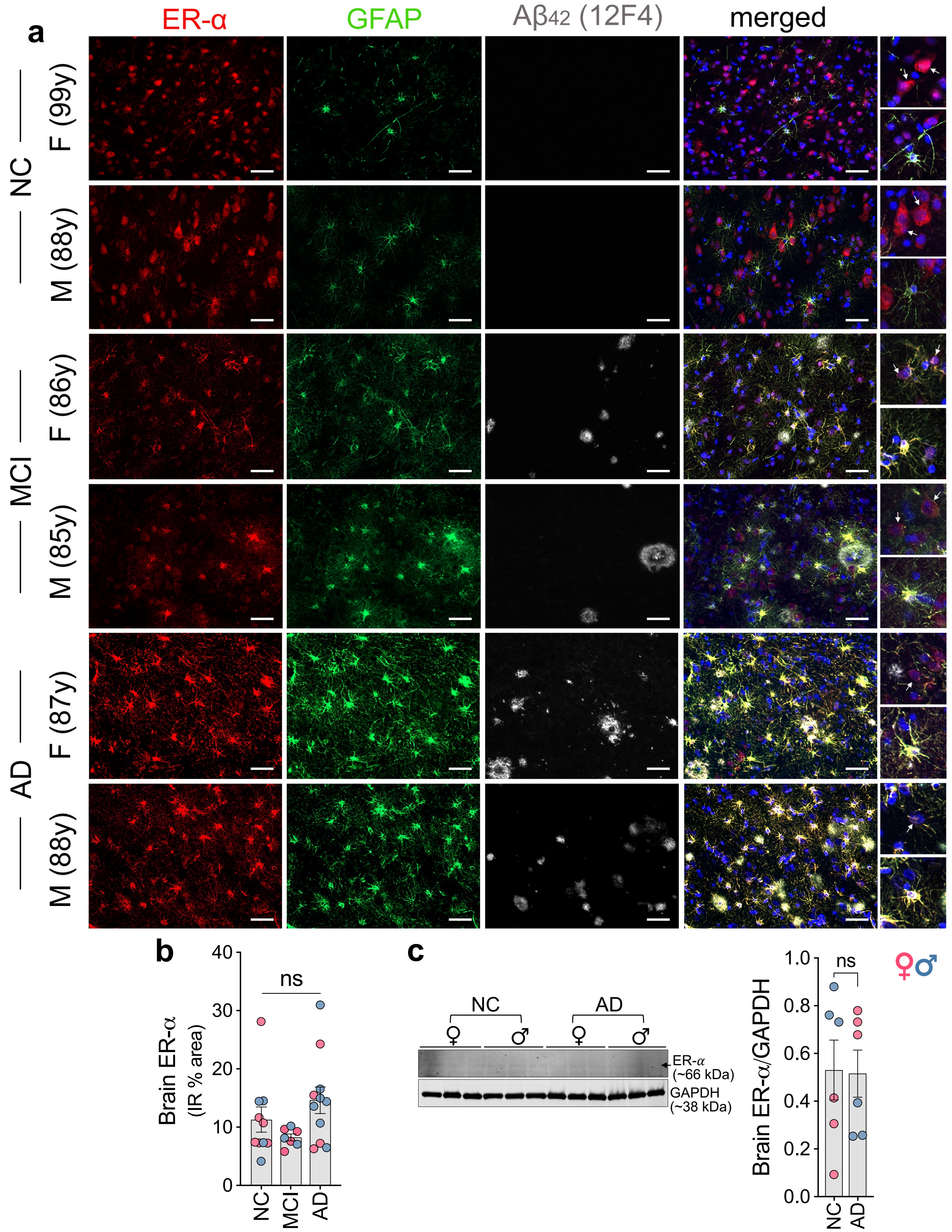


**Extended data Fig. 6: Expression of ER-α in the cerebral cortex of NC, MCI and AD individuals. a** Representative confocal images of postmortem cerebral cortex sections showing immunolabeling for the ER-α (red), astrocyte marker GFAP (green), and 12F4⁺ Aβ plaque in female and male MCI and AD patient versus NC control. Nuclei are stained with DAPI (blue). Scale bars: 20 µm. 2 repetitions. **b** Quantification of cortical ER-α⁺ immunoreactive (IR) % area across the diagnostic groups (NC=9, MCI=7, AD=10). **c** Representative immunoblots and densitometric analysis of ER-α in cerebral cortex of AD (n=6) versus NC donors (n=6). Data are represented as individual values (circles) and group means ± SEMs. 1-way ANOVA with Tukey's multiple comparison test for 3 or more groups and Student's 2-tailed *t*-test for 2-group comparisons. *ns: nonsignificant.*

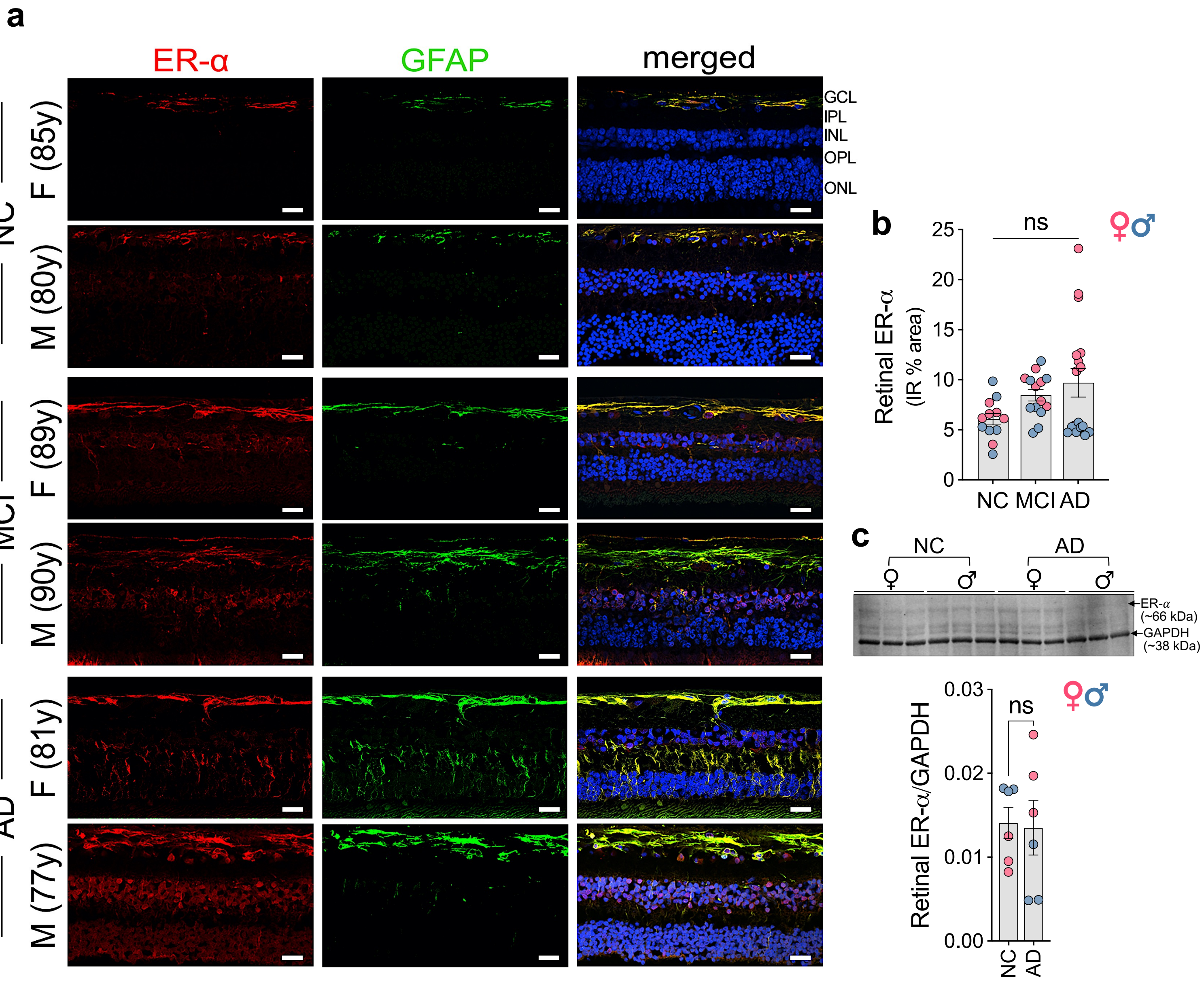


**Extended data Fig. 7: Expression of ER-α in the retina of NC, MCI and AD individuals: a** Representative confocal images showing immunolabelling for ER-α (red) and GFAP (green, astrocyte and Müller glia), in the postmortem retinal sections from female and male NC, MCI, and AD individuals. Nuclei are stained with DAPI (blue). Scale bars: 20 μm. 3 repetitions. NFL, neurofilament layer; GCL, ganglionic cell layer; INL, inner nuclear layer; OPL, ONL, outer nuclear layer. **b** Quantification of retinal ER-α$^{+}$ IR % areas across the diagnostic groups (NC=15, MCI=17, AD=17). **c** Representative immunoblots and densitometric analysis of ER-α in retina of AD (n=6) versus NC donors (n=6). Data are represented as individual values (circles) and group means ± SEMs. 1-way ANOVA with Tukey's multiple comparison test for 3 or more groups and Student's 2-tailed *t*-test for 2-group comparisons. *ns: nonsignificant.*

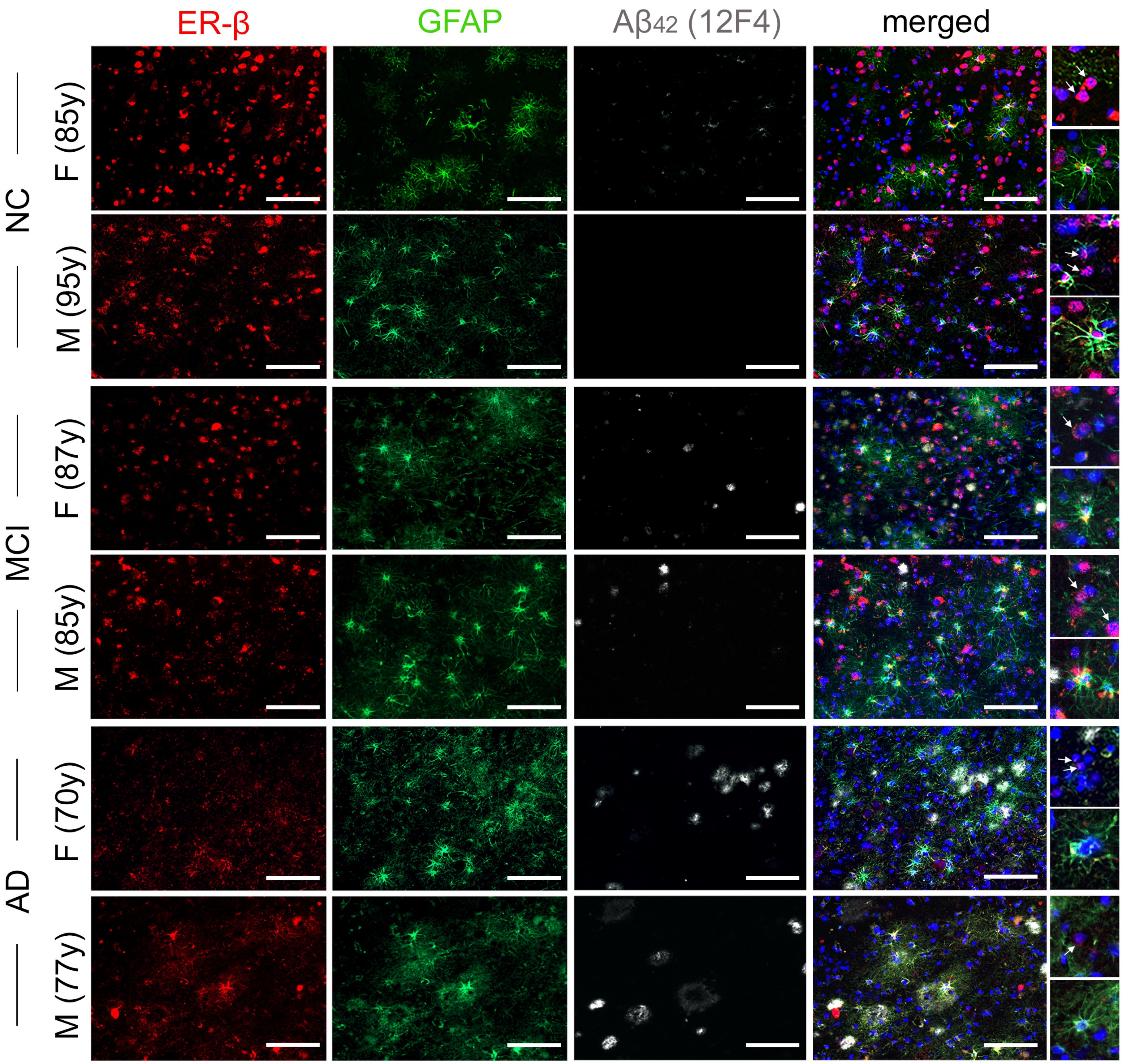


**Extended data Fig. 8: Expression of ER-β in the cerebral cortex of NC, MCI and AD individuals.** Representative confocal images of postmortem cerebral cortex sections showing immunolabeling for the ER-β (red), astrocyte marker GFAP (green), and 12F4$^{+}$ Aβ plaque in female and male MCI and AD patient versus NC control. Nuclei are stained with DAPI (blue). Scale bars: 20 µm. 2 repetitions.

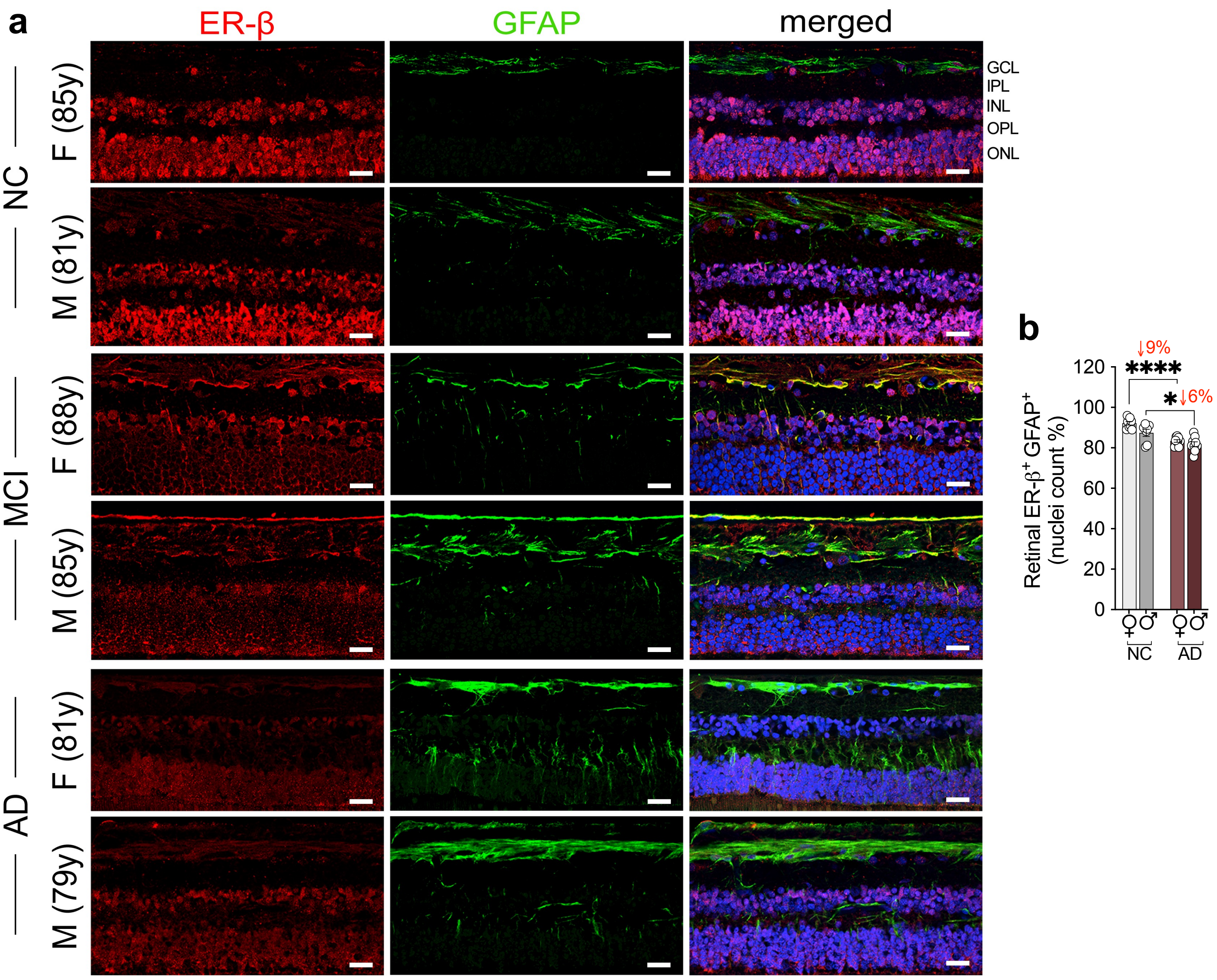


**Extended data Fig. 9: Expression of ER-β in the retina of NC, MCI and AD individuals. a** Representative confocal images of postmortem retinal sections showing immunolabeling for the ER-β (red) and GFAP (green, astrocyte and Müller glia) in female and male MCI and AD patient versus NC control. Nuclei are stained with DAPI (blue). Scale bars: 20 μm. 3 repetitions. NFL, neurofilament layer; GCL, ganglionic cell layer; INL, inner nuclear layer; OPL, ONL, outer nuclear layer. **b** Quantification of ER-β$^+$ GFAP$^+$ retinal astrocytic nuclei in female and male AD (n = 9F/9M) and NC (n = 9F/7M) retinas. Percentage change are indicated in red. Data are presented as individual subjects (circles), and group means ± SEMs. Statistical analyses used 2-way ANOVA with Bonferroni's post hoc tests. **P ≤ 0.05, ****P ≤ 0.0001.*

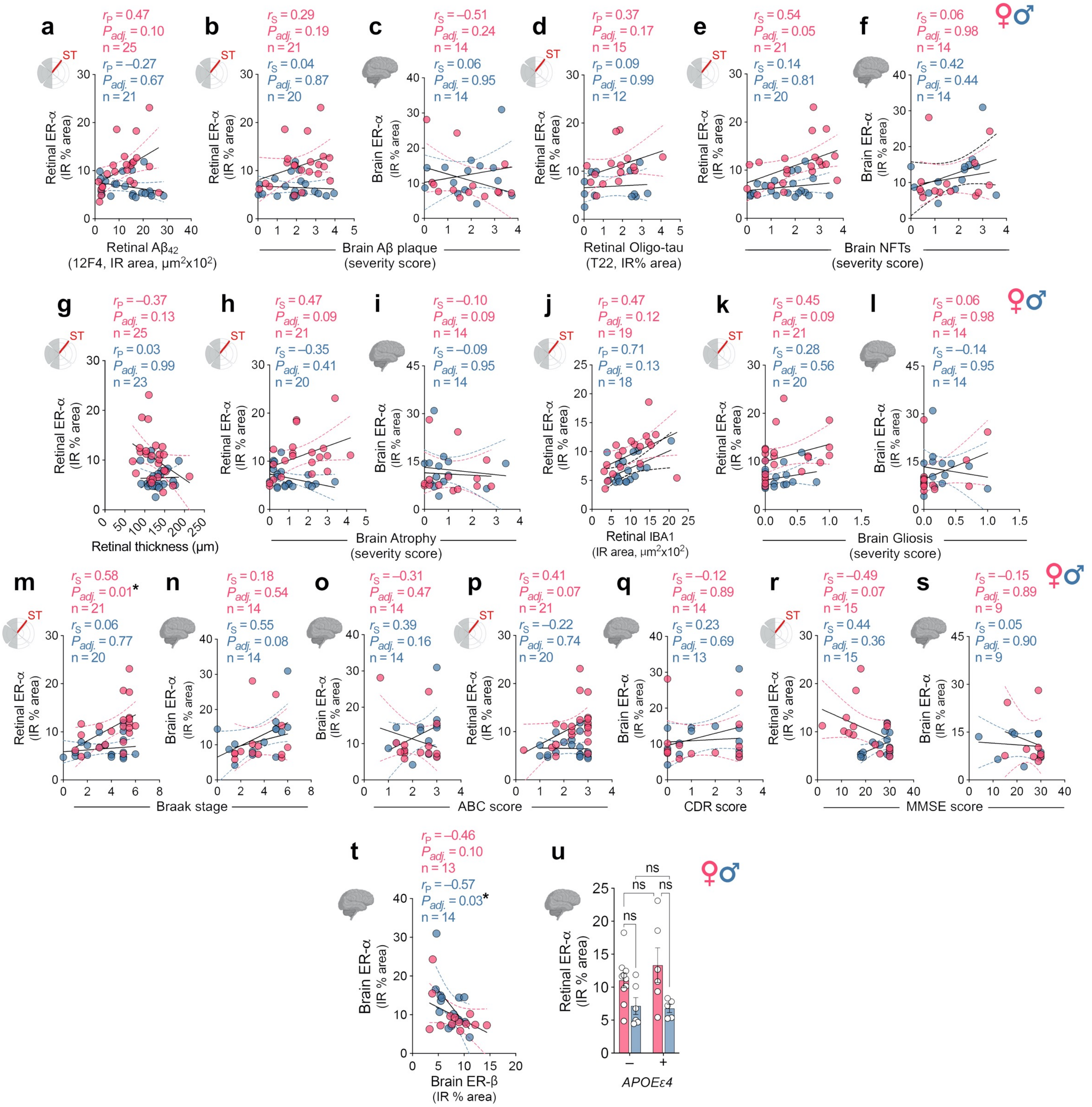


**Extended data Fig. 10: Sex-specific associations of retinal and cerebral ER-α with disease stage, cognition, and *APOEε4* genotype. a-t** Pearson and Spearman correlation between retinal or cerebral ER-α with **(a-c)** retinal and cerebral Aβ burden, **(d-f)** retinal and cerebral pathological tau burden, **(g-i)** retinal thickness and brain atrophy, **(j-l)** retinal and brain gliosis, **(m,n)** Braak stage, **(o,p)** ABC score, **(q)** CDR score, and **(r,s)** MMSE score. **(t)** Pearson correlation between brain ER-α and ER-β. **u** Sex-specific stratification of retinal ER-α across *APOEε4* carriers and non-carriers (n = 6F/9M ε4 carriers; 13F/9M, non-carriers). Data are presented as individual subjects (circles), and group means ± SEMs. 2-way ANOVA with Bonferroni's multiple comparison test for 3 or more group comparisons. For each correlation plot, Pearson or Spearman correlation coefficient (*r*), Holm-Šídák adjusted *P* values (asterisks), and number of individuals (n) are shown in the upper right corner of each graph. Black lines represent linear regression fits with 95% confidence interval. **P ≤ 0.05, ns: nonsignificant.*

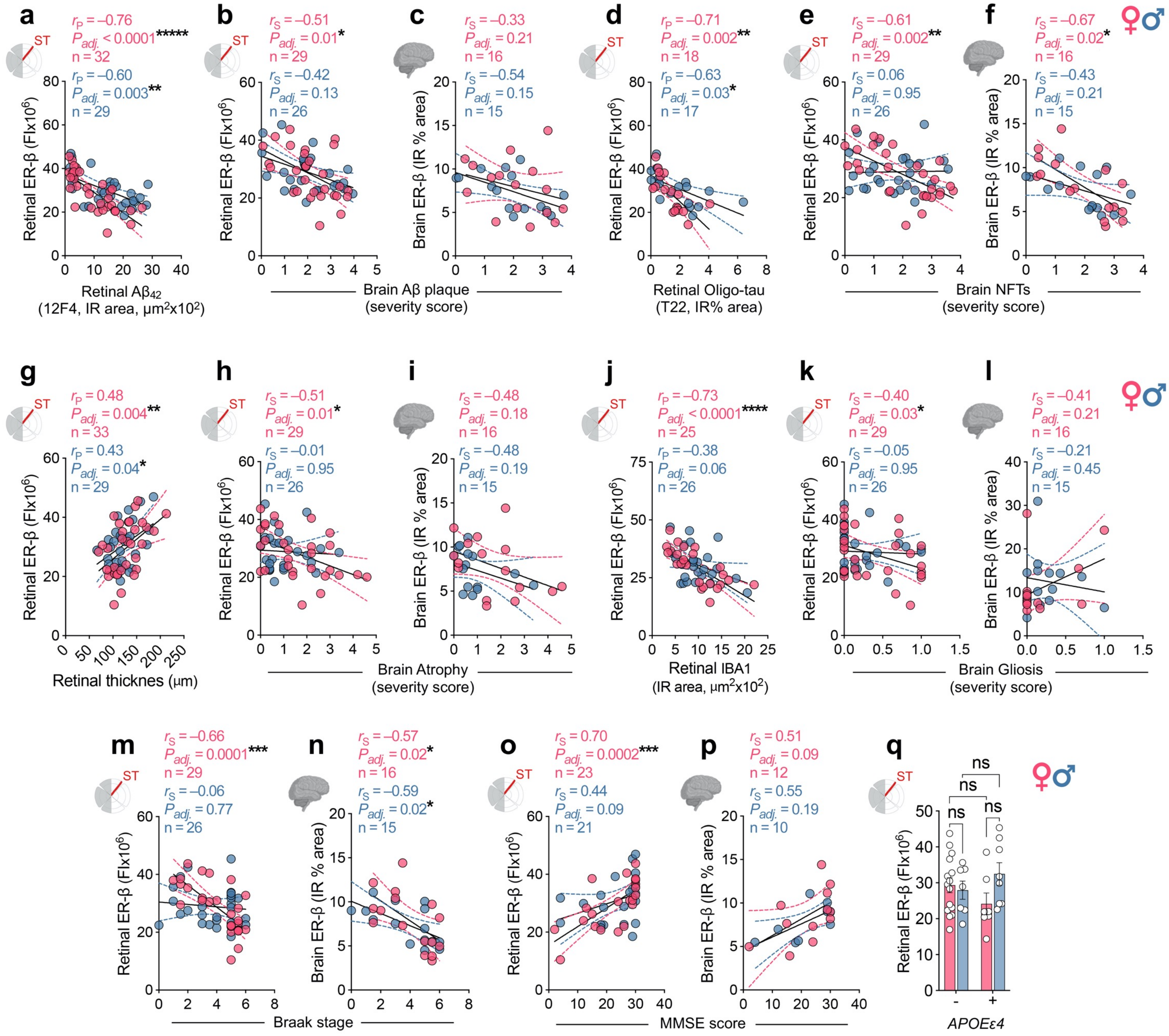


**Extended data Fig. 11: Sex-specific associations of retinal and cerebral ER-β with disease stage, cognition, and *APOEε4* genotype. a-p** Pearson and Spearman correlation analyses between retinal or cerebral ER-β and **(a-c)** retinal and cerebral Aβ burden, **(d-f)** retinal and cerebral pathological tau burden, **(g-i)** retinal thickness and brain atrophy, **(j-l)** retinal and brain gliosis, **(m,n)** Braak stage, and **(o,p)** MMSE score. **q** Sex-specific stratification of retinal ER-β across *APOEε4* carriers and non-carriers (n = 7F/8M ε4 carriers; 15F/7M, non-carriers). Data are presented as individual subjects (circles), and group means ± SEMs. 2-way ANOVA with Bonferroni's multiple comparison test for 3 or more group comparisons. For each correlation plot, Pearson or Spearman correlation coefficient (*r*), Holm-Šídák adjusted *P* values (asterisks), and number of individuals (n) are shown in the upper right corner of each graph. Black lines represent linear regression fits with 95% confidence interval. **P ≤ 0.05, **P ≤ 0.01, ***P ≤ 0.001, ****P ≤ 0.0001, ns: nonsignificant.*
